# A seventeenth-century *Mycobacterium tuberculosis* genome supports a Neolithic emergence of the *Mycobacterium tuberculosis* complex

**DOI:** 10.1101/588277

**Authors:** Susanna Sabin, Alexander Herbig, Åshild J. Vågene, Torbjörn Ahlström, Gracijela Bozovic, Caroline Arcini, Denise Kühnert, Kirsten I. Bos

## Abstract

**Background:** Although tuberculosis accounts for the highest mortality from a bacterial infection on a global scale, questions persist regarding its origin. One hypothesis based on modern *Mycobacterium tuberculosis* complex (MTBC) genomes suggests their most recent common ancestor (MRCA) followed human migrations out of Africa ~70,000 years before present (BP). However, studies using ancient genomes as calibration points have yielded much younger MRCA dates of less than 6,000 years. Here we aim to address this discrepancy through the analysis of the highest-coverage and highest quality ancient MTBC genome available to date, reconstructed from a calcified lung nodule of Bishop Peder Winstrup of Lund (b. 1605 – d. 1697).

**Results:** A metagenomic approach for taxonomic classification of whole DNA content permitted the identification of abundant DNA belonging to the human host and the MTBC, with few non-TB bacterial taxa comprising the background. Subsequent genomic enrichment enabled the reconstruction of a 141-fold coverage *M. tuberculosis* genome. In utilizing this high-quality, high-coverage 17^th^ century *M. tuberculosis* genome as a calibration point for dating the MTBC, we employed multiple Bayesian tree models, including birth-death models, which allowed us to model pathogen population dynamics and data sampling strategies more realistically than those based on the coalescent.

**Conclusions:** The results of our metagenomic analysis demonstrate the unique preservation environment calcified nodules provide for DNA. Importantly, we estimate an MRCA date for the MTBC of 3683 BP (2253-5821 BP) and for Lineage 4 of 1651 BP (946-2575 BP) using multiple models, confirming a Neolithic emergence for the MTBC.

## BACKGROUND

Tuberculosis, caused by organisms in the *Mycobacterium tuberculosis* complex (MTBC), has taken on renewed relevance and urgency in the 21^st^ century due to its global distribution, its high morbidity, and the rise of antibiotic resistant strains (1). The difficulty in disease management and treatment, combined with the massive reservoir the pathogen maintains in human populations through latent infection (2), makes tuberculosis a pressing public health challenge. Despite this, controversy exists regarding the history of the relationship between members of the MTBC and their human hosts.

Existing literature suggests two most recent common ancestor (MRCA) dates for the MTBC based on the application of Bayesian molecular dating to genome-wide *Mycobacterium tuberculosis* data. One estimate suggests the extant MTBC emerged through a bottleneck approximately 70,000 years ago, coincident with major migrations of humans out of Africa (3). This estimate was reached using exclusively modern *M. tuberculosis* genomes, with internal nodes of the MTBC calibrated by extrapolated dates for major human migrations (3). This estimate relied on congruence between the topology of MTBC and human mitochondrial phylogenies, but this congruence does not extend to human Y chromosome phylogeographic structure (4). As an alternative approach, the first publication of ancient MTBC genomes utilized radiocarbon dates as direct calibration points to infer mutation rates, and yielded an MRCA date for the complex of less than 6,000 years (5). This younger emergence was later supported by mutation rates estimated within the pervasive Lineage 4 (L4) of the MTBC, using four *M. tuberculosis* genomes from the late 18^th^ and early 19^th^ centuries (6).

Despite the agreement in studies that have relied on ancient DNA calibration so far, dating of the MTBC emergence remains controversial. Such a young age cannot account for purported detection of MTBC DNA in archaeological material that predates the MRCA estimate (e.g. Baker et al. 2015; Hershkovitz et al. 2008; Masson et al. 2013; Rothschild et al. 2001), the authenticity of which has been challenged (11). Furthermore, constancy in mutation rates of the MTBC has been challenged on account of observed rate variation in modern lineages, combined with the unquantified effects of latency (12). The ancient genomes presented by Bos and colleagues, though isolated from human remains, were most closely related to *Mycobacterium pinnipedii*, a lineage of the MTBC associated with infections in seals and sea lions today (5). Given our unfamiliarity with the demographic history of tuberculosis in sea mammal populations (13), identical substitution rates between the pinniped lineage and human-adapted lineages of the MTBC cannot be assumed. Additionally, the identification of true genetic changes in archaeological specimens can be difficult given the similarities between MTBC and environmental mycobacterial DNA from the depositional context (14). Though the ancient genomes published by Kay and colleagues belonged to human-adapted lineages of the MTBC, and the confounding environmental signals were significantly reduced by their funerary context in crypts, two of the four genomes used for molecular dating were derived from mixed-strain infections (6). By necessity, diversity derived in each genome would have to be ignored for them to be computationally distinguished (6). Though ancient DNA is a valuable tool for answering the question of when the MTBC emerged, the available ancient data remains sparse and subject to case-by-case challenges.

Here, we contribute to clarifying the timing of the emergence of the MTBC and L4 using multiple Bayesian models of varying complexity through the analysis of a high-coverage 17^th^ century *M. tuberculosis* genome extracted from a calcified lung nodule. Removed from naturally mummified remains, the nodule provided an excellent preservation environment for the pathogen, exhibiting minimal infiltration by exogenous bacteria. The nodule and surrounding lung tissue also showed exceptional preservation of host DNA, thus showing promise for this tissue type in ancient DNA investigations.

## RESULTS

### Pathogen identification

Computed tomography (CT) scans of the mummified remains of Bishop Peder Winstrup of Lund revealed a calcified granuloma a few millimeters (mm) in size in the collapsed right lung together with two ~5 mm calcifications in the right hilum (Figure 1). Primary tuberculosis causes parenchymal changes and ipsilateral hilar lymphadenopathy that is more common on the right side (15). Upon resolution it can leave a parenchymal scar, a small calcified granuloma (Ghon Focus), and calcified hilar nodes, which are together called a Ranke complex. In imaging this complex is suggestive of previous tuberculosis infection, although histoplasmosis can have the same appearance (16). Histoplasmosis, however, is very rare in Scandinavia and more often seen in other parts of the world (e.g. the Americas) (17). The imaging findings were therefore considered to result from previous primary tuberculosis. One of the calcified hilar nodes was extracted from the remains during video-assisted thoracoscopic surgery, guided by fluoroscopy. The extracted material was further subsampled for genetic analysis. DNA was extracted from the nodule and accompanying lung tissue using protocols optimized for the recovery of ancient, chemically degraded, fragmentary genetic material (18). The metagenomic library was shotgun sequenced to a depth of approximately 3.7 million reads.

**Figure 1.**
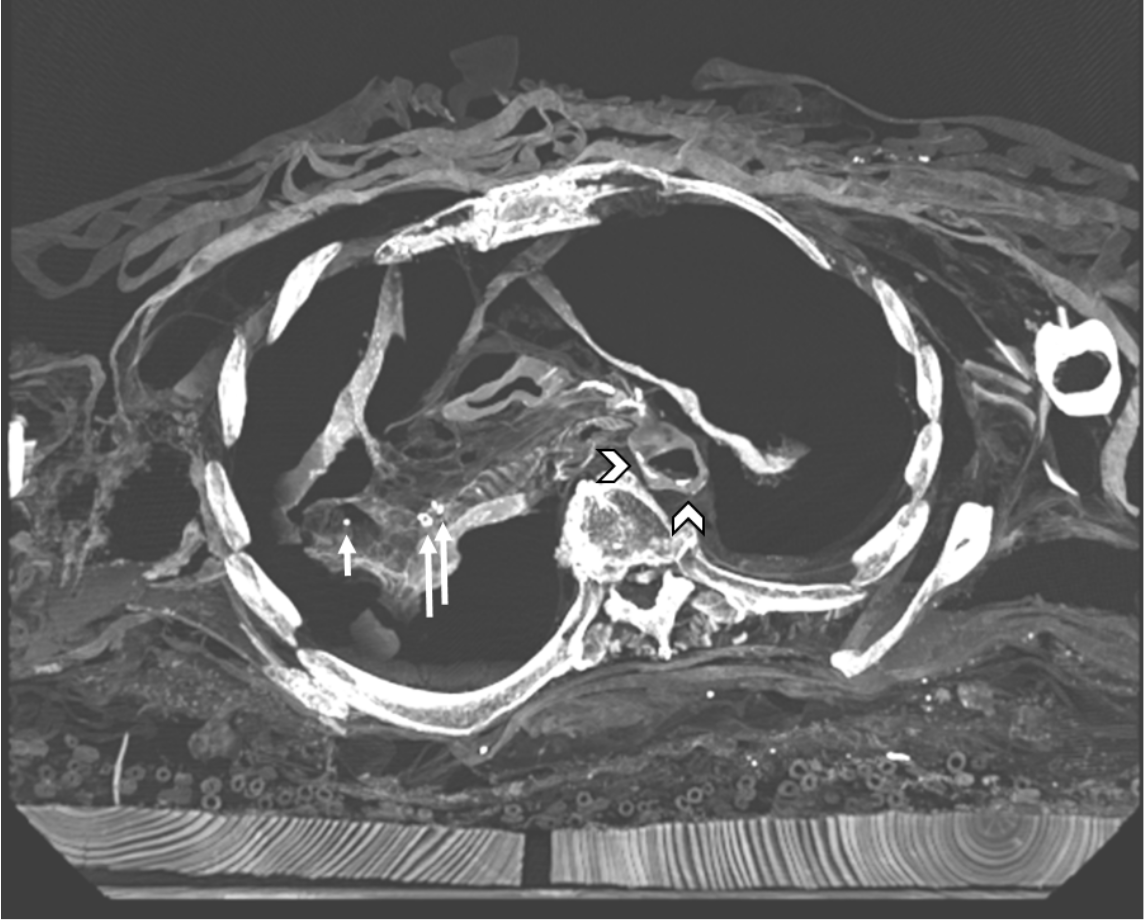
CT image of Ranke complex. CT image of Peder Winstrup’s chest in a slightly angled axial plane with the short arrow showing a small calcified granuloma in the probable upper lobe of the collapsed right lung, and two approximately 5 mm calcifications in the right hilum together suggesting a Ranke complex and previous primary tuberculosis. The more lateral of the two hilar calcifications was extracted for further analysis. In addition, there are calcifications in the descending aorta proposing atherosclerosis (arrowhead).

Adapter-clipped and base quality filtered reads were taxonomically binned with MALT (19) against the full NCBI Nucleotide database (‘nt’, April 2016). In this process, 3,515,715 reads, or 95% of the metagenomic reads, could be assigned to taxa contained within the database. Visual analysis of the metagenomic profile in MEGAN6 (20) revealed the majority of these reads, 2,833,403 or 81%, were assigned to *Homo sapiens*. A further 1,724 reads assigned to the *Mycobacterium tuberculosis* complex (MTBC) node. Importantly, no other taxa in the genus *Mycobacterium* were identified, and the only other identified bacterial taxon was *Ralstonia solanacearum* (Figure 2a), a soil-dwelling plant pathogen frequently identified in metagenomic profiles of archaeological samples (21,22) (Table S1 in Additional File 1).

**Figure 2.**
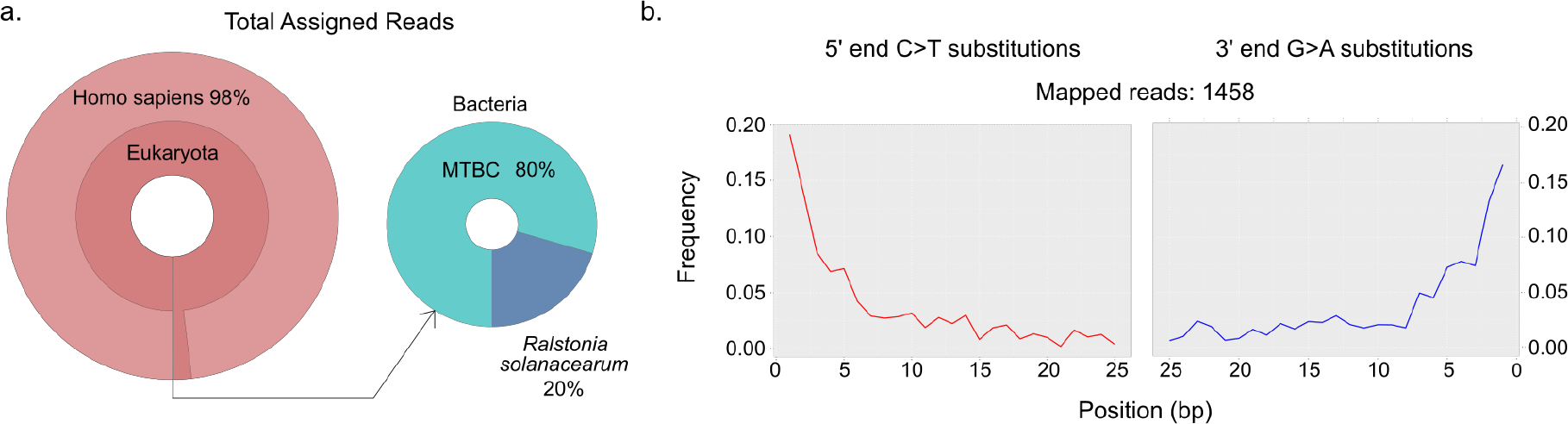
Screening of sequencing data from LUND1 shows preservation of host and pathogen DNA. A) Krona plots reflecting the metagenomic composition of the lung nodule. The majority of sequencing reads were aligned to *Homo sapiens* (n=2,833,403), demonstrating extensive preservation of host DNA. A small portion of reads aligned to bacterial organisms, and 80% of these reads were assigned to the MTBC node (n=1,724). B) Damage plots generated from sequencing reads mapped directly to a reconstructed MTBC ancestor genome (23), demonstrating a pattern characteristic of ancient DNA.

Pre-processed reads were mapped to both the hg19 human reference genome and a reconstructed MTBC ancestor (TB ancestor) (23) using BWA as implemented in the Efficient Ancient Genome Reconstruction (EAGER) pipeline (24). Reads aligned to hg19 with direct mapping constituted an impressive 88% of the total sequencing data (Table S2 in Additional File 1). Human mitochondrial contamination was extremely low, estimated at only 1-3% using Schmutzi (25) (Additional File 2). Reads were also mapped to the TB ancestor (Table 1). After map quality filtering and read de-duplication, 1,458 reads, or 0.045% of the total sequencing data, aligned to the reference (Table 1), and exhibited cytosine-to-thymine damage patterns indicative of authentic ancient DNA (Figure 2b) (26,27). Qualitative preservation of the tuberculosis DNA was slightly better than that of the human DNA, as the damage was greater in the latter (Table S2 in Additional File 1). Laboratory-based contamination, as monitored by negative controls during the extraction and library preparation processes, could be ruled out as the source of this DNA (Table S3 in Additional File 1).

**Table 1.** Mapping statistics for LUND1 libraries. A comparison of the mapping statistics for the non-UDG screening library and UDG-treated MTBC enriched library of LUND1 when aligned to the MTBC ancestor genome (23). For full EAGER output, see Table S2 in Additional File 1.

| Pre/post | Library treatment | Processed reads pre-mapping (*n*) | Unique mapped reads, quality-filtered (*n*) | Endogenous DNA (%) | Mean fold coverage | Mean fragment length (bp) | GC content (%) |
| --- | --- | --- | --- | --- | --- | --- | --- |
| **Pre-capture** | non-UDG | 3696712 | 1458 | 0.045 | 0.018 | 54.31 | 63.89 |
| **Post-capture** | UDG | 59091507 | 9482901 | 45.652 | 141.5062 | 65.83 | 62.96 |

### Genomic enrichment and reconstruction

Due to the clear but low-abundance MTBC signal, a uracil DNA glycosylase (UDG) library was constructed to remove DNA lesions caused by hydrolytic deamination of cytosine residues (28) and enriched with an in-solution capture (29,30) designed to target genome-wide data representing the full diversity of the MTBC (see METHODS). The capture probes are based on the TB ancestor genome (23), which is equidistant from all lineages of the MTBC. The enriched library was sequenced using a paired-end, 150-cycle Illumina sequencing kit to obtain a full fragment-length distribution (Figure S1 in Additional File 2). The resulting sequencing data was then aligned to the hypothetical TB ancestor genome (23), and the mapping statistics were compared with those from the screening data to assess enrichment (Table 1). Enrichment increased the proportion of endogenous MTBC DNA content by three orders of magnitude, from 0.045% to 45.652%, and deep sequencing yielded genome-wide data at an average coverage of approximately 141.5-fold. The mapped reads have an average fragment length of ~66 base pairs (Table 1).

We further evaluated the quality of the reconstructed genome by quantifying the amount of heterozygous positions (see METHODS). Derived alleles represented by 10-90% of the reads covering a given position with five or more reads of coverage were counted. Only 24 heterozygous sites were counted across all variant positions in LUND1. As a comparison, the other high-coverage (~125 fold) ancient genome included here – body92 from Kay et al. 2015 – contained 70 heterozygous positions.

### Phylogeny and dating

Preliminary phylogenetic analysis using neighbor joining (Figures S2 and S3 in Additional File 2), maximum likelihood (Figures S4 and S5 in Additional File 2), and maximum parsimony trees (Figures S6 and S7 in Additional File 2) indicated that LUND1 groups within the L4 strain diversity of the MTBC, and more specifically, within the L4.10/PGG3 sublineage. This sublineage was recently defined by Stucki and colleagues as the clade containing L4.7, L4.8, and L4.9 (31) according to the widely-accepted Coll nomenclature (32). Following this, we constructed two datasets to support molecular dating of the full MTBC (Table S4 in Additional File 1) and L4 of the MTBC (Table S5 in Additional File 1).

The dataset reflecting extant diversity of the MTBC was compiled as reported elsewhere (5), with six ancient genomes as calibration points. These included LUND1; two additional ancient genomes, body80 and body92, extracted from late 18^th^ and early 19^th^ century Hungarian mummies (6); and three human-isolated *Mycobacterium pinnipedii* strains from Peru (5), encompassing all available ancient *M. tuberculosis* genomes with sufficient coverage to call SNPs confidently after stringent mapping with BWA (33) (see METHODS; Table S4 in Additional File 1). *Mycobacterium canettii* was used as an outgroup. In generating an alignment of variant positions in this dataset, we excluded repetitive regions and regions at risk of cross-mapping with other organisms as done previously (5), as well as potentially imported sites from recombination events, which were identified using ClonalFrameML (34) (Table S6 in Additional File 1). We chose to exclude these potential recombination events despite *M. tuberculosis* being generally recognized as a largely clonal organism with minimal recombination or horizontal gene transfer, as this is still a point of contention (35). Only twenty-three variant sites were lost from the full MTBC alignment as potential imports. We called a total of 42,856 variable positions in the dataset as aligned to the TB ancestor genome. After incompletely represented sites were excluded, 11,716 were carried forward for downstream analysis.

To explore the impact of the selected tree prior and clock model, we ran multiple variations of models as available for use in BEAST2 (36). We first used both a strict and a relaxed clock model together with a constant coalescent model (CC+strict, CC+UCLD). We found there to be minimal difference between the inferred rates estimated by the two models. This finding, in addition to the low rate variance estimated in all models, suggests there is little rate variation between known branches of the MTBC. Nevertheless, the relaxed clock appeared to have a slightly better performance (Table 2). To experiment with models that allowed for dynamic populations, we applied a Bayesian skyline (SKY+UCLD) and birth-death skyline prior (BDSKY+UCLD) combined with a relaxed clock model. In the BDSKY+UCLD model, the tree was conditioned on the root. To our knowledge, this is the first instance of a birth-death tree prior being used to infer evolutionary dynamics of the MTBC while using ancient data for tip calibration.

**Table 2.** Model comparison for full MTBC dataset. Parameter estimates from four models applied to the full MTBC dataset: constant coalescent with uncorrelated lognormal clock (CC+UCLD), constant coalescent with strict clock (CC+strict), Bayesian skyline coalescent with uncorrelated lognormal clock (SKY+UCLD), and birth-death skyline with uncorrelated lognormal clock (BDSKY+UCLD).

| Model | Mean Likelihood | Mean Rate (95% HPD) | Mean Rate Variance (95% HPD) | Mean Tree Height (95% HPD) |
| --- | --- | --- | --- | --- |
| BDSKY+UCLD | -6123180.475 | 1.303E-8<br>(6.9753E-9, 1.8348E-8) | 1.3656E-17<br>(2.4838E-18, 2.5884E-17) | 3683.203<br>(2253.2836, 5820.8405) |
| CC+UCLD | -6123187.492 | 1.214E-8<br>(7.1934E-9, 1.6448E-8) | 1.2459E-17<br>(2.833E-18, 2.3969E-17) | 4172.1961<br>(2585.2349, 6119.744) |
| SKY+UCLD | -6123279.053 | 1.3294E-8<br>(8.9335E-9, 1.7461E-8) | 1.4147E-17<br>(5.1837E-18, 2.4356E-17) | 3540.7193<br>(2453.8322, 4829.7259) |
| CC+strict | -6123688.933 | 1.1573E-8<br>(8.6397E-9, 1.4509E-8) | NA | 4453.1162<br>(3330.1516, 5619.3974) |

A calibrated maximum clade credibility (MCC) tree was generated for the BDSKY+UCLD model, with 3683 years before present (BP) (95% highest posterior density [95% HPD] interval: 2253 – 5821 BP) as an estimated date of emergence for the MTBC (Figure 3a). Tree topology agrees with previously presented phylogenetic analyses of the full MTBC (3,5,37). A birth-death skyline plot illustrates the flux in the effective reproduction number (R) over time (Figure 3b). In an outbreak setting, R refers to the average number of secondary cases stemming from a single infection, and an epidemic event is inferred when the value is greater than one. However, for the data at hand, R > 1 translates to lineage diversification rates exceeding lineage death/extinction. Since there is no data representing the period between ~1000 years ago and the emergence of the MTBC, there is much uncertainty in the related estimates. From around 1300 BP the 95% HPD excludes 1, indicating a positive net diversification rate, with a significant increase between 974 and 390 BP (odds ratio=10.00054).

**Figure 3.**
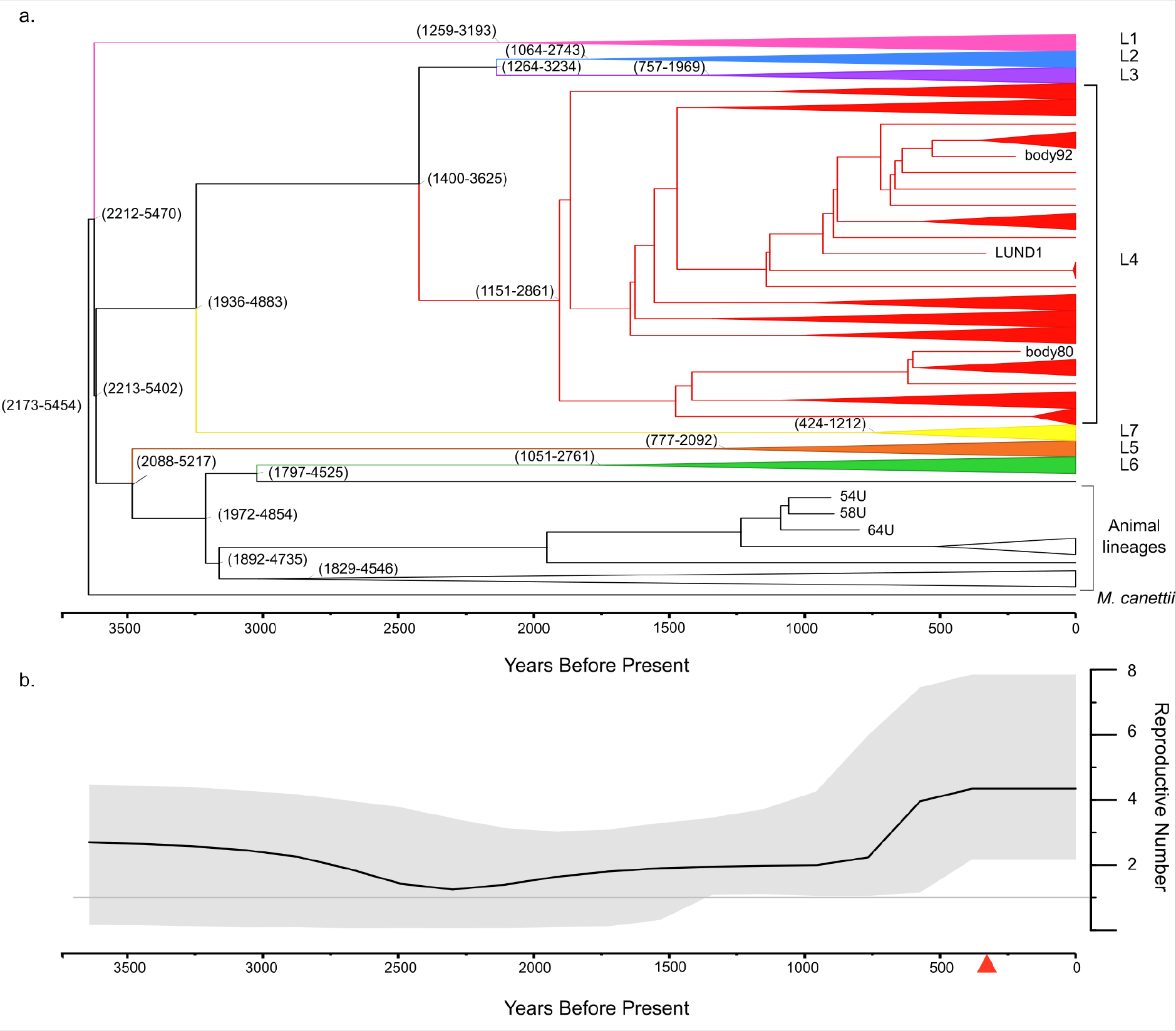
MTBC maximum clade credibility tree and birth-death skyline plot. A) This MCC tree of mean heights was generated from the BDSKY+UCLD model as applied to the full MTBC dataset. Modern genomes are collapsed according to lineage (labeled on the right side). The ancient genomes are labeled with their sample name. The outgroup is labeled as “*M. canettii*.” The 95% HPD intervals of selected node heights are indicated as (lower boundary -upper boundary) in years before present. The time scale is expressed as years before present, with the most recent time as 2010. B) The black line indicates median reproductive number over time (reproductive number set to 5 dimensions, see Additional File 2). The shaded grey area represents the 95% HPD interval of the reproductive number. The grey line indicates a reproductive number of 1. The red triangle on the timeline indicates the temporal position of LUND1.

The L4 dataset includes LUND1 and the two Hungarian mummies described above (6) as calibration points. We selected 149 modern genomes representative of the known diversity of L4 from previously published datasets (Additional File 2) (3,23,31). A modern Lineage 2 (L2) genome was used as an outgroup. After the exclusion of sites as discussed above (Table S7 in Additional File 1), a SNP alignment of these genomes in reference to the reconstructed TB ancestor genome (23) included a total of 17,333 variant positions, excluding positions unique to the L2 outgroup. Only fifteen variant sites were lost from the L4 dataset alignment. After sites missing from any alignment in the dataset were excluded from downstream analysis, 10,009 SNPs remained for phylogenetic inference. A total of 810 SNPs were identified in LUND1, of which 126 were unique to this genome. A SNP effect analysis (38) was subsequently performed on these derived positions (Additional File 2; Table S8 in Additional File 1).

We applied the same models as described above for the full MTBC dataset, with the addition of a birth-death skyline model conditioned on the origin of the root (BDSKY+UCLD+origin). All mean tree heights are within 250 years of each other and the 95% HPD intervals largely overlap. As an informal model comparison, the BDSKY+UCLD model shows the highest marginal likelihood values. We employed the BDSKY+UCLD+origin model to determine if the estimated origin of the L4 dataset agreed with the tree height estimates for the full MTBC dataset. Intriguingly, the estimated origin parameter (Table 3), or the ancestor of the tree root, largely overlaps with the 95% HPD range for MTBC tree height as seen in Table 2.

**Table 3.** Model comparison for L4 dataset. Selected parameter estimates from five models applied to the Lineage 4 dataset: constant coalescent with uncorrelated lognormal clock (CC+UCLD), constant coalescent with strict clock (CC+strict), Bayesian skyline coalescent with uncorrelated lognormal clock (SKY+UCLD), birth-death skyline with uncorrelated lognormal clock and tree conditioned on the root (BDSKY+UCLD), and birth-death skyline with uncorrelated lognormal clock with origin parameter estimate (BDSKY+UCLD+origin).

| Model | Mean Likelihood | Mean Rate (95% HPD) | Mean Rate Variance (95% HPD) | Mean Tree Height (95% HPD) | Origin (BDSKY only) |
| --- | --- | --- | --- | --- | --- |
| BDSKY+UCLD | -6032794.215 | 2.8369E-8<br>(1.4945E-8, 4.0535E-8) | 4.071E-17<br>(5.4068E-18, 7.7999E-17) | 1650.608<br>(945.7849, 2574.5831) | NA |
| BDSKY+UCLD+<br>origin | -6032794.987 | 3.1477E-8<br>(2.0046E-8, 4.2189E-8) | 4.5822E-17<br>(9.6022E-18, 8.051E-17) | 1462.2611<br>(935.6968, 2058.31) | 3268.341<br>(1102.2144, 8071.0277) |
| CC+UCLD | -6032797.480 | 3.1068E-8<br>(1.988E-8, 4.1624E-8) | 4.3865E-17<br>(1.3291E-17, 7.806E-17) | 1569.0512<br>(1054.607, 2225.4758) | NA |
| SKY+UCLD | -6032874.480 | 2.8097E-8<br>(1.5329E-8, 3.9927E-8) | 3.7609E-17<br>(6.0593E-18, 7.1919E-17) | 1690.536<br>(1016.2712, 2646.5163) | NA |
| CC+strict | -6033002.535 | 2.9299E-8<br>(2.2173E-8, 3.6637E-8) | NA | 1567.544<br>(1186.1186, 1978.6488) | NA |

A calibrated MCC tree (Figure 4a) was generated based on the BDSKY+UCLD model for the L4 dataset. This model yielded an estimated date of emergence for L4 of 1650 BP (95% HPD: 946-2575 BP). The tree reflects the ten-sublineage topology presented by Stucki and colleagues (31), with LUND1 grouping with the L4.10/PGG3 sublineage. A birth-death skyline plot was also generated (Figure 4b), which is similar to that generated for the MTBC (Figure 3b) inasmuch as the mean R was continuously greater than one in the L4 population since its emergence.

**Figure 4.**
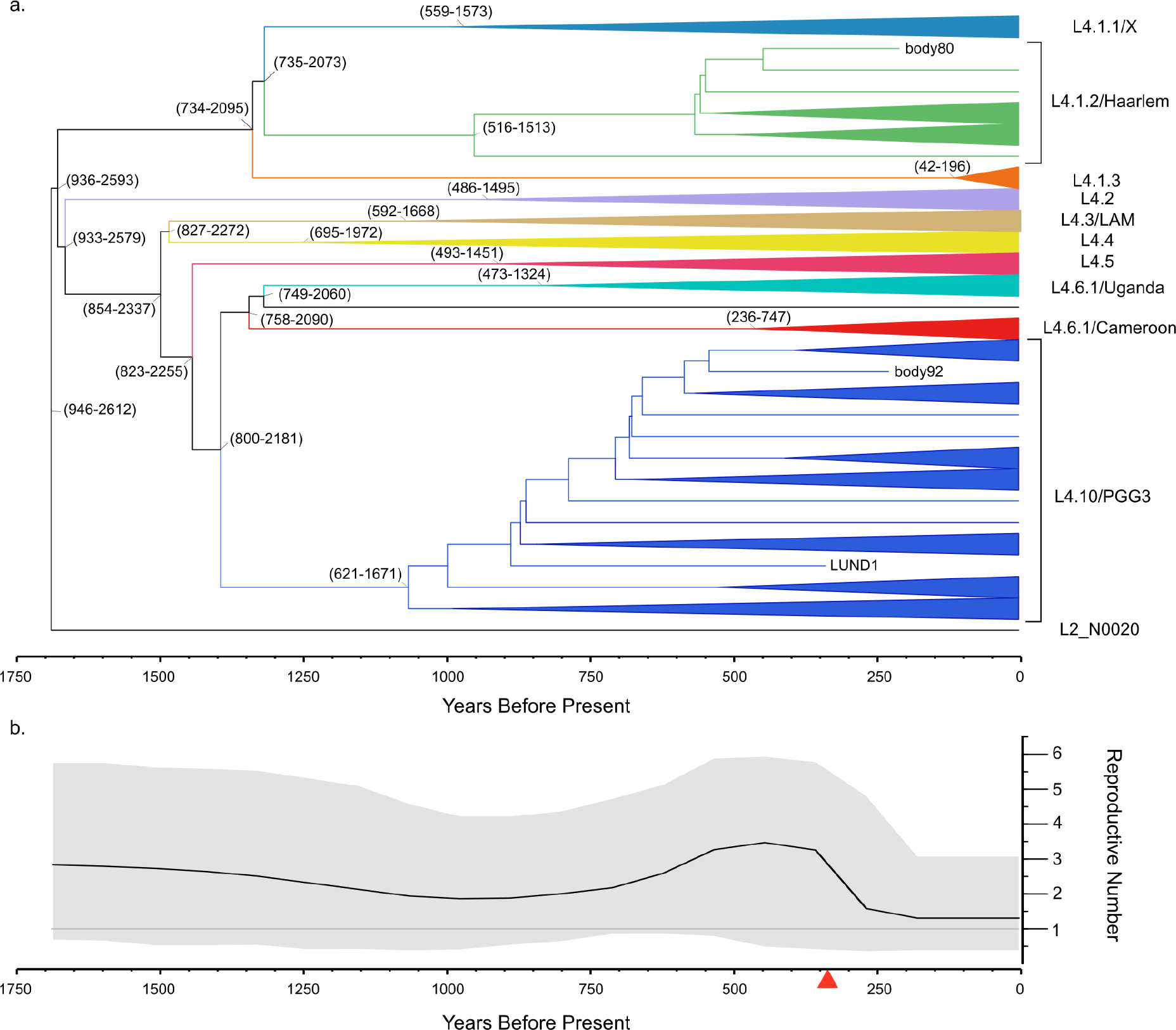
L4 maximum clade credibility tree and birth-death skyline plot. A) This MCC tree of mean heights was generated from the BDSKY+UCLD model as applied to the L4 dataset. Modern genomes are collapsed according to sublineage (labeled on the right side). The ancient genomes are labeled with their sample name. The Lineage 2 outgroup is labeled as “L2_N0020.” The 95% HPD interval of selected node heights is indicated as (lower boundary-upper boundary) in years before present. The time scale is expressed as years before present, with the most recent time as 2010. B) The black line indicates median reproductive number over time (reproductive number set to 5 dimensions, see Additional File 2). The shaded grey area represents the 95% HPD interval of the reproductive number. The grey line indicates a reproductive number of 1. The red triangle on the timeline indicates the position of LUND1.

## DISCUSSION

The increasing number of ancient *Mycobacterium tuberculosis* genomes is steadily reducing the uncertainty of molecular dating estimates for the emergence of the MTBC. Here, using the ancient data available to date, we directly calibrate the MTBC time tree, and confirm that known diversity within the complex is derived from a common ancestor that existed ~2000-6000 years before present (Figure 3; Table 2) (5,6). Our results support the hypothesis that the MTBC emerged during the Neolithic, and not before. The Neolithic revolution generally refers to the worldwide transition in lifestyle and subsistence from more mobile, foraging economies to more sedentary, agricultural economies made possible by the domestication of plants and animals. The period during which it occurred varies between regions. In Africa, where the MTBC is thought to have originated (3,39–41), the spread of animal domestication in the form of pastoralism appears to have its focus around ~3000 BCE, or 5000 BP, across multiple regions (42). The estimates presented here place the emergence of tuberculosis amidst the suite of human health impacts that took place as a consequence of the Neolithic lifestyle changes often referred to collectively as the first epidemiological transition (43,44).

Tuberculosis has left testaments to its history as a human pathogen in the archaeological record (45), and some skeletal evidence has implied the existence of tuberculosis in humans and animals pre-dating the lower 95% HPD boundary for the MTBC MRCA presented here (7,8,10,46–50). However, it is important to explore the evolutionary history of the MTBC through molecular data. Furthermore, it is crucial to base molecular dating estimates on datasets that include ancient genomes, which expand the temporal sampling window and provide data from the pre-antibiotic era. Numerous studies have found long-term nucleotide substitution rate estimates in eukaryotes and viruses to be dependent on the temporal breadth of the sampling window, and it is reasonable to assume the same principle applies to bacteria (51–56). Additionally, rate variation over time and between lineages, which may arise due to changing evolutionary dynamics such as climate and host biology, can impact the constancy of the molecular clock (54,55). Though models have been developed to accommodate uncertainty regarding these dynamics (57), temporally structured populations can provide evidence and context for these phenomena over time and can aid researchers in refining models appropriate for the taxon in question (56).

In addition to our MRCA estimate for the MTBC, we present one for L4, which is among the most globally dominant lineages in the complex (31,58). Our analyses yielded MRCA dates between ~1000-2500 years before present, as extrapolated from the 95% HPD intervals of all models (Table 3), with the mean dates spanning from 320-548 CE. These results are strikingly similar to those found in two prior publications, and support the idea proposed by Kay and colleagues that L4 may have emerged during the late Roman period (5,6). However, there exist discrepancies between different estimates for the age of this lineage in available literature that touch the upper (37) and lower (58) edges of the 95% HPD intervals reported here. In addition, recent phylogeographic analyses of the MTBC and its lineages had ambiguous results for L4, with the internal nodes being assigned to either African or European origins depending on the study or different dataset structures used within the same study (37,58). Despite the ambiguity, this finding belies a close relationship between ancestral L4 strains in Europe and Africa (37,58). Stucki and colleagues delineated L4 into globally distributed “generalist” sublineages and highly local “specialist” sublineages that do not appear outside a restricted geographical niche (31). Thus far, the specialist sublineages are limited to the African continent; however, a clear phylogenetic relationship explaining the distinction between geographically expansive and limited strains has not been established. Specifically, LUND1 falls within the globally distributed, “generalist” L4.10/PGG3 sublineage that shares a clade with two specialist sublineages: L4.6.1/Uganda and L4.6.2/Cameroon (Figure 4) (31). In the BDSKY+UCLD model presented here, the ancestral node for this clade dates to approximately 1372 years BP. At this extrapolated time, an ancestral strain underwent an evolutionary event in which some descendant lineages acquired or lost a feature that equipped them to expand past limited host niches into Eurasia. Confirming and elucidating this phenomenon could offer relevant clues regarding the evolutionary relationship between populations of MTBC organisms and humans. However, the current discrepancies over the age and geographic origin of L4 make interpretations of existing data unreliable for this purpose. These discrepancies could be due to differences in genome selection, SNP selection, and/or model selection and parameterization. Until more diverse, high-quality ancient L4 genomes are generated, creating a more temporally and geographically structured dataset, it is unlikely we will gain clarity.

Going deeper into comparisons between the results presented here and those from prior studies, mutation rate estimates in the L4 and full MTBC analyses were lower than previous estimates for comparable datasets, but within the same order of magnitude, with all mean and median estimates ranging between 1E-8 and 5E-8 (5,6) (Table 2). Nucleotide substitution rates inferred based on modern tuberculosis data are close to, but slightly higher than those based on ancient calibration, with multiple studies finding rates of approximately 1E-7 substitutions per site per year in multiple studies (4,59). Despite a strict clock model having been rejected by the MEGA-CC molecular clock test (60) for both the L4 and full MTBC datasets, the clock rate variation estimates do not surpass 9E-17 in any model. Additionally, there is little difference between the clock rates estimated in the L4 and full MTBC datasets suggesting the rate of evolution in L4 does not meaningfully differ from that of the full complex (Tables 2 and 3; Figure 5; Figures S9 and S10 in Additional File 2).

**Figure 5.**
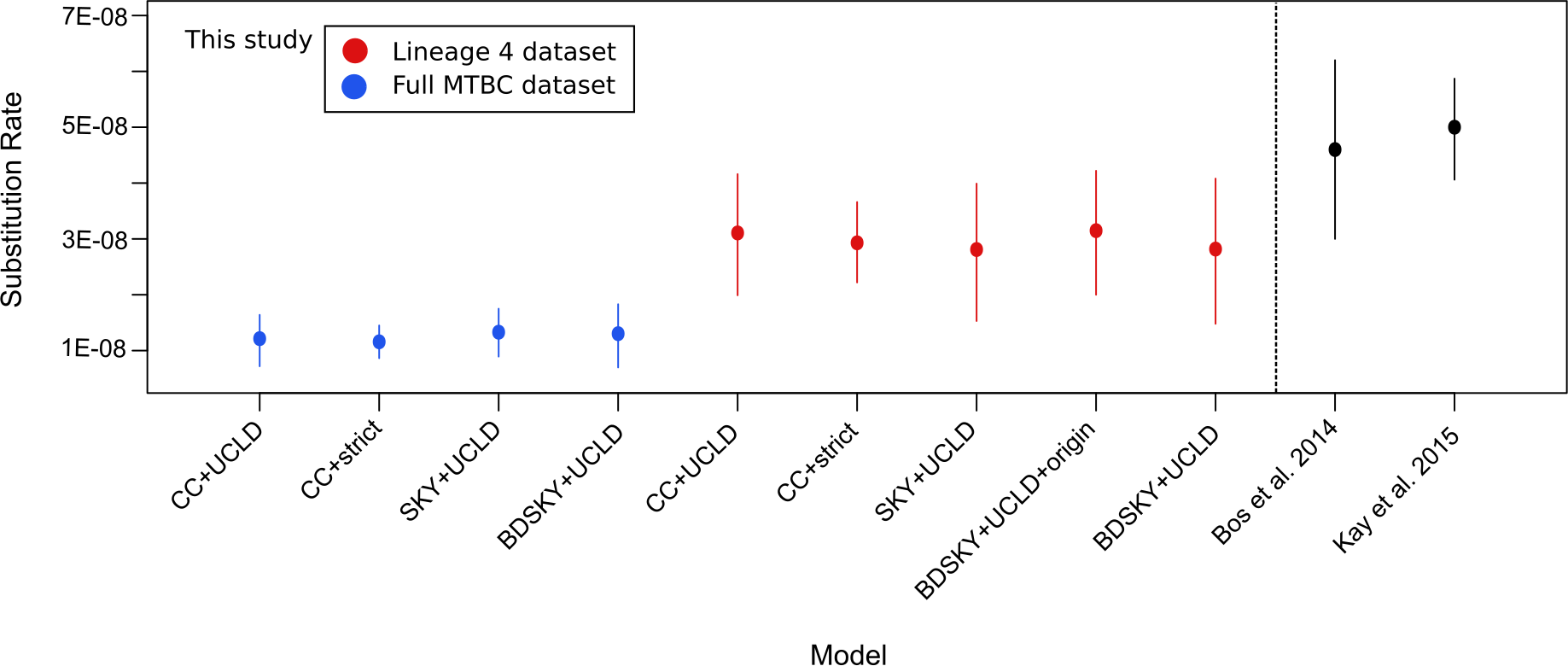
Substitution rate comparison across models and studies. Mean substitution rate per site per year for all models is expressed by a filled circle, with extended lines indicating the 95% HPD interval for that parameter. The Bos et al. 2014 and Kay et al. 2015 ranges are based on the reported rate values in each study. The Bos et al. 2014 range is based on a full MTBC dataset, while the Kay et al. 2015 range is based on an L4 dataset. All values presented here fall within one order of magnitude.

Another parameter explored here is R over time for the MTBC and L4 (Figures 2b and 3b). For both datasets, we see an increase in R at approximately 750 BP. In the MTBC model, it increases sharply and maintains its peak between 4 and 5. The increase is more gradual in the L4 model, and declines to hovering just above R=1. This roughly coincides with a jump in effective population size estimated by Liu and colleagues for MTBC lineages indigenous to China (61). The decline of R for L4 beginning approximately 350 BP appears surprising, given the historically recorded rise of the White Plague in Europe from the 17^th^-19^th^ century (62). However, this is likely due to the reduced sampling of modern sequences, which enabled the dating of the entire L4 lineage. A thorough phylodynamic analysis requires inclusion of “outbreak samples” only (63) and shall be explored in future work.

Importantly, we explored our data through multiple models, including birth-death tree priors. In our opinion, these models offer more robust parameterization options for heterochronous datasets that are unevenly distributed over time, such as those presented here, by allowing for uneven sampling proportions across different time intervals of the tree (64). Recent studies have demonstrated the importance of selecting appropriate tree priors for the population under investigation, as well as the differences between birth-death and coalescent tree priors (65,66). It is notable that the estimates reported here roughly agree across multiple demographic and clock models implemented in BEAST2. The estimate of the origin height for the L4 dataset as calculated with the birth-death Skyline model overlaps with the 95% HPD intervals for the tree height estimates across models in the full MTBC dataset.

In addition to confirming the findings of prior publications, this study contributes a high-coverage, contamination-free, and securely dated ancient *M. tuberculosis* genome for future dating efforts, which may include more ancient data or more realistic models. Much of this quality likely comes from the unique preservation environment of the calcified nodule. In the case of tuberculosis, such nodules form from host immunological responses in the waning period of an active pulmonary infection and remain in lung tissue, characterizing the latent form of the disease. Host immune cells were likely responsible for the dominant signal of 89% human DNA in the LUND1 metagenomic screening library. Similar levels of preservation have been observed through analyses of ancient nodules yielding *Brucella* (Kay et al, 2014) and urogenital bacterial infections (DeVault et al, 2017), with pathogen preservation rivaling what we report here.

LUND1 avoided multiple quality-related problems often encountered in the identification and reconstruction of ancient genetic data from the MTBC. The genome is of high quality both in terms of its high coverage and low heterozygosity. Despite the low quantity of MTBC DNA detected in the preliminary screening data, in-solution capture enriched the proportion of endogenous DNA by three orders of magnitude (Table 1). The resultant genomic coverage left few ambiguous positions at which multiple alleles were represented by greater than 10% of the aligned reads. This extremely low level of heterozygosity indicated that LUND1 contained a dominant signal of only one MTBC strain. This circumvented analytical complications that can arise from the simultaneous presence of multiple MTBC strains associated with mixed infections, or from the presence of abundant non-MTBC mycobacteria stemming from the environment. The preservation conditions of Bishop Winstrup’s remains, mummified in a crypt far from soil, left the small MTBC signal unobscured by environmental mycobacteria or by the dominance of any other bacterial organisms (Figure 2a). The unprecedented quality of LUND1 and the precision of its calibration point (historically recorded year of death) made it ideal for Bayesian molecular dating applications.

As the practice of applying ancient genomic data is still in its nascent stages, there are caveats to the results of this study. First, this analysis excludes *M. canettii* – a bacterium that can cause pulmonary tuberculosis – from the MTBC dataset, and as such our estimate does not preclude the possibility of a closely related ancestor having caused tuberculosis-like infections in humans before 6,000 BP. The inferred MRCA could be restricted to a lineage that survived an evolutionary bottleneck, possibly connected to its virulence in humans as suggested elsewhere, albeit as a considerably more ancient event (67,68). Additionally, despite the use of ancient data, our temporal sampling window is still narrow given the estimated age of the MTBC and L4. For the MTBC dataset no samples pre-date 1,000 years before present, and for L4, no samples predate 350 years before present. It could be argued the ancient L4 genomes available to date represent samples taken in the midst of an epidemic – namely, the “White Plague” of tuberculosis, which afflicted Europe between the 17^th^ and 19^th^ centuries (62). For a slow-evolving bacterial pathogen like tuberculosis, it is possible our sampling window of ancient genomes is subject to the very issue they are meant to alleviate: the time-dependency of molecular clocks (51,53–55). The genomes sampled from pre-contact Peruvian remains do not derive from a known epidemic period in history and add temporal spread to our MTBC dataset. However, their membership to a clade of animal-associated strains (*M. pinnipedii*) indicates they were subject to dramatically different evolutionary pressures compared to the human-associated lineages of the complex due to differing host biology and population dynamics. On a related matter, the available ancient MTBC genomes also suffer from a lack of lineage diversity, with only pinniped strains and L4 represented.

Filling the MTBC time tree with more ancient genomes from diverse time periods, locations, and lineages would address the limitations listed above. The most informative data would a) derive from an Old World context (i.e. Europe, Asia, or Africa) pre-dating the White Plague in Europe or b) come from any geographical location or pre-modern time period, but belong to one of the MTBC lineages not yet represented by ancient data. An ideal data point, which would clarify many open questions and seeming contradictions related to the evolutionary history of the MTBC, would derive from Africa, the inferred home of the MTBC ancestor (3,39–41), and pre-date 2,000 years before present. A genome of this age would test the lower boundaries of the 95% HPD tree height intervals estimated in the full MTBC models presented here. Until recently it would have been considered unrealistic to expect such data to be generated from that time period and location. Innovations and improvements in ancient DNA retrieval and enrichment methods, however, have brought this expectation firmly into the realm of the possible (30,69). Ancient bacterial pathogen genomes have now been retrieved from remains from up to 5,000 years before present (70–72) and recent studies have reported the recovery of human genomes from up to 15,000 year-old remains from north Africa (73,74).

## CONCLUSIONS

Here we offer confirmation that the extant MTBC, and all available ancient MTBC genomes, stem from a common ancestor that existed a maximum of 6,000 years before present. Many open questions remain, however, regarding the evolutionary history of the MTBC and its constituent lineages, as well as the role of tuberculosis in human history. Elucidating these questions is an iterative process, and progress will include the generation of diverse ancient *M. tuberculosis* genomes, and the refinement and improved parameterization of Bayesian models that reflect the realities of MTBC (and other organisms’) population dynamics and sampling frequencies over time. To aid in future attempts to answer these questions, this study provides an ancient MTBC genome of impeccable quality and explores the first steps in applying birth-death population models to modern and ancient TB data.

## METHODS

### Lung nodule identification

The paleopathological investigation of the body of Winstrup is based on extensive CT-scan examinations with imaging of the mummy and its bedding performed with a Siemens Somatom Definition Flash, 128 slice at the Imaging Department of Lund University Hospital. Ocular inspection of the body other than of the head and hands was not feasible, since Winstrup was buried in his episcopal robes and underneath the body was wrapped in linen strips. The velvet cap and the leather gloves were removed during the investigation. The body was naturally mummified and appeared to be well preserved with several internal organs identified.

The imaging was quite revealing. The intracranial content was lost with remains of the brain in the posterior skull base. Further, the dental status was poor with several teeth in the upper jaw affected by severe attrition, caries and signs of tooth decay, as well as the absence of all teeth in the lower jaw. Most of the shed teeth were represented by closed alveoli, indicating antemortem tooth loss. Along with the investigation of the bedding, a small sack made of fabric was found behind the right elbow containing five teeth: two incisors, two premolars and one molar. The teeth in the bag complemented the remaining teeth in the upper jaw. It is feasible that the teeth belonged to Winstrup and were shed several years before he died. A fetus approximately five months of age was also found in the bedding, underneath his feet.

Both lungs were preserved but collapsed with findings of a small parenchymal calcification and two ~5 mm calcifications in the right hilum (Figure 1). The assessment was that these could constitute a Ranke complex, suggestive of previous primary tuberculosis. A laparoscopy was performed at the Lund University Hospital in a clinical environment whereby the nodules were retrieved. Furthermore, several calcifications were also found in the aorta and the coronary arteries, suggesting the presence of atherosclerosis. The stomach, liver and gall bladder were preserved, and several small gallstones were observed. The spleen could be identified but not the kidneys. The intestines were there, however, collapsed except for the rectum that contained several large pieces of concernments. The bladder and the prostate could not be recognized.

The skeleton showed several pathological changes. Findings on the vertebrae consistent with of DISH (Diffuse idiopathic skeletal hyperostosis) were present in the thoracic and the lumbar spine. Reduction of the joint space in both hip joints and the left knee joint indicate that Winstrup was affected by osteoarthritis. No signs of gout or osteological tuberculosis (i.e. Pott’s disease) were found.

Neither written sources nor the modern examination of the body of Winstrup reveal the immediate cause of death. However, it is known that he was bedridden for at least two years preceding his death. Historical records indicate that gallstones caused him problems while travelling to his different parishes. Additionally, he was known to have suffered from tuberculosis as a child, which may have recurred in his old age.

### Sampling and extraction

Sampling of the lung nodule, extraction, and library preparation were conducted in dedicated ancient DNA clean rooms at the Max Planck Institute for the Science of Human History in Jena, Germany. The nodule was broken using a hammer, and a 5.5 mg portion of the nodule was taken with lung tissue for extraction according to a previously described protocol with modifications (18). The sample was first decalcified overnight at room temperature in 1 mL of 0.5 M EDTA. The sample was then spun down, and the EDTA supernatant was removed and frozen. The partially decalcified nodule was then immersed in 1 mL of a digestion buffer with final concentrations of 0.45 M EDTA and 0.25 mg/mL Proteinase K (Qiagen) and rotated at 37°C overnight. After incubation, the sample was centrifuged. The supernatants from the digestion and initial decalcification step were purified using a 5 M guanidine-hydrochloride binding buffer with a High Pure Viral Nucleic Acid Large Volume kit (Roche). The extract was eluted in 100 μl of a 10mM tris-hydrochloride, 1 mM EDTA (pH 8.0), and 0.05% Tween-20 buffer (TET). Two negative controls and one positive control sample of cave bear bone powder were processed alongside LUND1 to control for reagent/laboratory contamination and process efficiency, respectively.

### Library preparation and shotgun screening sequencing

Double-stranded Illumina libraries were constructed according to an established protocol with some modifications (75). Overhangs of DNA fragments were blunt-end repaired in a 50 μl reaction including 10 μl of the LUND1 extract, 21.6 μl of H_2_O, 5 μl of NEB Buffer 2 (New England Biolabs), 2 μl dNTP mix (2.5 mM), 4 μl BSA (10 mg/ml), 5 μl ATP (10 mM), 2 μl T4 polynucleotide kinase, and 0.4 μl T4 polymerase, then purified and eluted in 18 μl TET. Illumina adapters were ligated to the blunt-end fragments in a reaction with 20 μl Quick Ligase Buffer, 1 μl of adapter mix (0.25 μM), and 1 μl of Quick Ligase. Purification of the blunt-end repair and adapter ligation steps was performed using MinElute columns (Qiagen). Adapter fill-in was performed in a 40 μl reaction including 20 μl adapter ligation eluate, 12 μl H_2_O, 4 μl Thermopol buffer, 2 μl dNTP mix (2.5 mM), and 2 μl Bst polymerase. After the reaction was incubated at 37°C for 20 minutes, the enzyme was heat deactivated with a 20 minute incubation at 80°C. Four library blanks were processed alongside LUND1 to control for reagent/laboratory contamination. The library was quantified using a real-time qPCR assay (Lightcycler 480 Roche) with the universal Illumina adapter sequences IS7 and IS8 as targets. Following this step, the library was double indexed (76) with a unique pair of indices over two 100 μl reactions using 19 μl of template, 63.5 μl of H_2_O, 10 μl PfuTurbo buffer, 1 μl PfuTurbo (Agilent), 1 μl dNTP mix (25mM), 1.5 μl BSA (10 mg/ml), and 2 μl of each indexing primer (10 μM). The master mix was prepared in a pre-PCR clean room and transported to a separate lab for amplification. The two reactions were purified and eluted in 25 μl of TET each over MinElute columns (Qiagen), then assessed for efficiency using a real-time qPCR assay targeting the IS5 and IS6 sequences in the indexing primers. The reactions were then pooled into one double-indexed library. Approximately one-third of the library was amplified over three 70 μl PCR reactions using 5 μl of template each and Herculase II Fusion DNA Polymerase (Agilent). The products were MinElute purified, pooled, and quantified using an Agilent Tape Station D1000 Screen Tape kit. LUND1 and the corresponding negative controls were sequenced separately on an Illumina NextSeq 500 using single-end, 75-cycle, high-output kits.

### Pathogen identification and authentication

De-multiplexed sequencing reads belonging to LUND1 were processed *in silico* with the EAGER pipeline (v.1.92) (24). ClipAndMerge was used for adapter removal, fragment length filtering (minimum sequence length: 30 bp), and base sequence quality filtering (minimum base quality: 20). MALT v. 038 (19) was used to screen the metagenomic data for pathogens using the full NCBI Nucleotide database (‘nt’, April 2016) with a minimum percent identity of 85%, a minSupport threshold of 0.01, and a topPercent value of 1.0. The resulting metagenomic profile was visually assessed with MEGAN6 CE (20). The adapter-clipped reads were additionally aligned to a reconstructed MTBC ancestor genome (23) with BWA (33) as implemented in EAGER (-l 1000, -n 0.01, -q 30). Damage was characterized with DamageProfiler in EAGER (77).

### In-solution capture probe design

Single-stranded probes for in-solution capture were designed using a computationally extrapolated ancestral genome of the MTBC (23). The probes are 52 nucleotides in length with a tiling density of 5 nucleotides, yielding a set of 852,164 unique probes after the removal of duplicate and low complexity probes. The number of probes was raised to 980,000 by a random sampling among the generated probe sequences. A linker sequence (5’-CACTGCGG-3’) was attached to each probe sequence, resulting in probes of 60 nucleotides in length, which were printed on a custom-design 1 million-feature array (Agilent). The printed probes were cleaved off the array, biotinylated and prepared for capture according to Fu et al. (30).

### UDG library preparation and in-solution capture

Fifty microliters of the original LUND1 extract were used to create a uracil-DNA glycosylase (UDG) treated library, in which the post-mortem cytosine to uracil modifications, which cause characteristic damage patterns in ancient DNA, are removed. The template DNA was treated in a buffer including 7 μl H_2_O, 10 μl NEB Buffer 2 (New England Biolabs), 12 μl dNTP mix (2.5 mM), 1 μl BSA (10 mg/ml), 10 μl ATP (10 mM), 4 μl T4 polynucleotide kinase, and 6 μl USER enzyme (New England Biolabs). The reaction was incubated at 37°C for three hours, then 4 μl of T4 polymerase was added to the library to complete the blunt-end repair step. The remainder of the library preparation protocol, including double indexing, was performed as described above.

The LUND1 UDG-treated library was amplified over two rounds of amplification using Herculase II Fusion DNA Polymerase (Agilent). In the first round, five reactions using 3 μl of template each were MinElute purified and pooled together. The second round of amplification consisted of three reactions using 3 μl of template each from the first amplification pool. The resulting products were MinElute purified and pooled together. The final concentration of 279 ng/μl was measured using an Agilent Tape Station D1000 Screen Tape kit (Agilent). A portion of the non-UDG library (see above) was re-amplified to 215 ng/μl. A 1:10 pool of the non-UDG and UDG amplification products was made to undergo capture. A pool of all associated negative control libraries (Supplementary Table 2) and a positive control known to contain *M. tuberculosis* DNA also underwent capture in parallel with the LUND1 libraries. Capture was performed according to an established protocol (29), and the sample product was sequenced on an Illumina HiSeq 4000 with a 150-cycle paired end kit to a depth of ~60 million paired reads. The blanks were sequenced on a NextSeq 500 with a 75-cycle paired end kit.

### Genomic reconstruction, heterozygosity, and SNP calling

For the enriched, UDG-treated LUND1 sequencing data, de-multiplexed paired-end reads were processed with the EAGER pipeline (v. 1.92) (24), adapter-clipped with AdapterRemoval, and aligned to the MTBC reconstructed ancestor genome with in-pipeline BWA (-l 32, -n 0.1, -q 37). Previously published ancient and modern *Mycobacterium tuberculosis* genomic data (Supplementary Table 4, Supplementary Table 5) were processed as single-end sequencing reads, but otherwise processed identically in the EAGER pipeline. Genome Analysis Toolkit (GATK) UnifiedGenotyper was used to call SNPs using default parameters and the EMIT ALL SITES output option (78). We used MultiVCFAnalyzer (v0.87 https://github.com/alexherbig/MultiVCFAnalyzer) (5) to create and curate SNP alignments for the L4 (Supplementary Table 5) and full MTBC (Supplementary Table 4) datasets based on SNPs called in reference to the TB ancestor genome (23), with repetitive sequences, regions subject to cross-species mapping, and potentially imported sites excluded. The repetitive and possibly cross-mapped regions were excluded as described previously (5). Potentially imported sites were identified using ClonalFrameML (34) separately for each dataset, using full genomic alignments and trees generated in RAxML (79) as input without the respective outgroups. Remaining variants were called as homozygous if they were covered by at least 5 reads, had a minimum genotyping quality of 30, and constituted at least 90% of the alleles present at the site. Outgroups for each dataset were included in the SNP alignments, but no variants unique to the selected outgroup genomes were included. Minority alleles constituting over 10% were called and assessed for LUND1 to check for a multiple strain *M. tuberculosis* infection. Sites with missing or incomplete data were excluded from further analysis.

### Phylogenetic analysis

Maximum likelihood, maximum parsimony, and neighbor joining trees were generated for the L4 and full MTBC datasets (Tables S4 and S5 in Additional File 1), with 500 bootstrap replications per tree. Maximum parsimony and neighbor joining trees were configured using MEGA-Proto and executed using MEGA-CC (60). Maximum likelihood trees were configured and executed using RAxML (79) with the GTR+GAMMA substitution model.

### Bayesian phylogenetic analysis of full MTBC and L4 datasets

Bayesian phylogenetic analysis of the full MTBC was conducted using a dataset of 261 *M. tuberculosis* genomes including LUND1, five previously published ancient genomes (5,6), and 255 previously published modern genomes (Table S4 in Additional File 1). *Mycobacterium canettii* was used as an outgroup for this dataset. Bayesian phylogenetic analysis of L4 of the MTBC was conducted using a dataset of 152 genomes including three ancient genomes presented here and in a previous publication (6) and 149 previously published modern genomes (Table S5 in Additional File 1). Body80 and body92 were selected out of the eight samples presented by Kay and colleagues based on multiple criteria. Multiple samples from that study proved to be mixed strain infections. Apart from body92, these samples were excluded from this analysis due to our present inability to separate strains without ignoring derived positions. Body92 had a clearly dominant strain estimated by Kay et al. (6) to make up 96% of the tuberculosis data, and stringent mapping in BWA (33) (-l 32, -n 0.1, -q 37) found the genome to have 124-fold coverage when mapped against the TB ancestor. Between the degree of dominance and the high coverage, we could confidently call variant positions from the dominant strain (Figure S8a in Additional File 2). Body80 was the only single-strain sample from that collection to have sufficient coverage (~8x) for confident SNP calling after stringent mapping (Figure S8b in Additional File 2). For selection criteria for the modern genomes, please see Additional File 2. L2_N0020 was used as an outgroup. The possibility of equal evolutionary rates in both datasets was rejected by the MEGA-CC molecular clock test (60). TempEst (80) was also used to assess temporal structure in the phylogeny prior to analysis with BEAST2 (36) (full MTBC R^2^=0.273; L4 R^2^=0.113).

A correction for static positions in the *M. tuberculosis* genome not included in the SNP alignment was included in the configuration file. A “TVM” substitution model, selected based on results from ModelGenerator (81), was implemented in BEAUti as a GTR+G4 model with the AG rate parameter fixed to 1.0. LUND1, body80, and body92 were tip-calibrated using year of death, which was available for all three individuals (Table S5 in Additional File 1). The three ancient Peruvian genomes were calibrated using the mid-point of their OxCal ranges (Table S4 in Additional File 1) (5). We performed tip sampling for all modern genomes excluding the outgroup over a uniform distribution between 1992 and 2010 for all but the BDSKY models for both datasets. The outgroup was fixed to 2010 in every case. In the BDSKY models, all modern genomes were given a tip date of 2010. All tree priors were used in conjunction with an uncorrelated relaxed lognormal clock model. The constant coalescent model was also used in conjunction with a strict clock model.

Two independent MCMC chains of 200,000,000 iterations minimum were computed for each model. If the ESS for any parameter was below 200 after the chains were combined, they were resumed with additional iterations. The results were assessed in Tracer v1.7.1 with a 10 percent burn-in (82). Trees were sampled every 20,000 iterations. The log files and trees for each pair of runs were combined using LogCombiner v2.4.7 (36). An MCC tree was generated using TreeAnnotator with ten percent burn-in (36). For details on the parameterization of the birth-death models, please see Additional File 2.

## DECLARATIONS

### Ethics approval and consent to participate

Not applicable

### Consent for publication

Not applicable

### Availability of data and material

Raw sequencing data from the non-UDG, non-enriched screening library and the UDG-treated, enriched library can be found under the BioProject PRJNA517266.

### Authors’ contributions

C.A. and K.I.B. conceived of the investigation. S.S., D.K., A.H., and K.I.B. designed the experiments. T.A., G.B., and C.A. performed the exhumation and radiological analysis of the mummy and provided a paleopathological examination. G.B. was responsible for the CT examinations together with imaging analysis and coordination of the calcification extraction. S.S. performed laboratory work. S.S., D.K., A.H., Å.J.V., and K.I.B. performed analyses.

## Acknowledgments

The authors would like to thank Marta Burri for conducting the hybridization capture. We also thank Elizabeth A. Nelson for laboratory assistance, Maria Spyrou and Felix M. Key technical assistance, and all members of the Molecular Palaeopathology and Computational Pathogenomics research groups at the MPI-SHH for constructive comments throughout the study. We would like to acknowledge the Department of Imaging and Clinical Physiology, Skåne University Hospital Lund for the generous opportunity to examine the mummy of Peder Winstrup and the radiologists Roger Siemund, Pär Wingren, Mats Geijer, Pernilla Gustavson and David Pellby for contributing to the image analyses. A sincere thank you to the thoracic surgeons Erik Gyllstedt and Jesper Andreasson for so delicately extracting the small calcification of interest and enabling further investigations.

## ADDITIONAL FILES

### Additional File 1

Format: Excel spreadsheet (.xlsx)

Title: Supplementary Tables

Description: Large tables of data contributing to the analyses presented in this paper, including a taxon table showing assigned reads from all taxonomic levels represented in the metagenomic LUND1 library (Table S1); full EAGER pipeline results for LUND1 shotgun sequencing data when mapped to HG19 human reference genome and TB ancestor genome, the non-UDG-treated enriched LUND1 data when mapped to the TB ancestor genome, and the UDG-treated enriched LUND1 data when mapped to the TB ancestor genome (Table S2); full EAGER pipeline results for negative controls processed with LUND1, mapped to the reconstructed TB ancestor genome (Table S3); genomes included in the full MTBC dataset, with respective publications, accession numbers, lineages, and dates (when applicable) (Table S4); genomes included in the L4 dataset, with respective publications, accession numbers, lineages, dates, and percentage of total SNPs called as heterozygous (Table S5); sites excluded from the full MTBC dataset (Table S6); sites excluded from the L4 dataset (Table S7); SnpEff annotation for derived alleles in LUND1 (Table S8).

### Additional File 2

Format: Word document (.docx)

Title: Supplementary Information

Description: Detailed supplements to the RESULTS and METHODS sections, including supplementary figures.

