## Additional File 2 for "A seventeenth-century *Mycobacterium tuberculosis* genome supports a Neolithic emergence of the *Mycobacterium tuberculosis* complex"

For

**Mitochondrial analysis**

Given the uncharacteristically high endogenous human DNA content of the lung nodule, we estimated the mitochondrial contamination rate using Schmutzi (Renaud et al. 2015). In order to do this we mapped the metagenomic sequencing data to the hg19 mitochondrial chromosome with CircularMapper as implemented in the EAGER pipeline (Peltzer et al. 2016). We called initial contamination estimates with contDeam, and then ran Schmutzi, calling a consensus mitochondrial sequence and making a final average contamination estimate of 2% (lower boundary 1%, upper boundary 3%).

**Heterozygosity analysis and selection of modern L4 genomes**

We chose to reduce the quantity of L4 genomes used for Bayesian phylogeny in this study for the purposes of computational feasibility and attempting to balance the representation of different L4 sublineages without compromising allelic diversity in the analysis. The four deeply sampled sublineages published by Stucki and colleagues (2016) – L4.3/LAM, L4.6.1/Uganda, L4.1.2/Haarlem, and L4.10/PGG3 – were subsampled to 30 representatives each based on the quantity of heterozygous positions found in each genome after the inclusion of strains published in the Comas et al. (2013, 2010) datasets. For each of these four sublineages, the 30 genomes with the fewest heterozygous positions were selected for inclusion. We quantified the number of heterozygous positions using MultiVCFAnalyzer (v0.87 https://github.com/alexherbig/MultiVCFAnalyzer) (Bos et al. 2014) (see METHODS).

**SNP Effect Analysis**

SnpEff (v4.2) (Cingolani et al. 2012) was run for LUND1 using a gene annotation database tailored to the TB ancestor reference genome (Comas et al. 2010). With the exception of setting the upstream and downstream effect interval to 100 nucleotides, we used default settings. Variant positions unique to LUND1 in comparison to the L4 dataset alignment were filtered, and the SnpEff results for these positions, in addition to gene descriptions from the NCBI Gene database, can be found in Table S9 of Additional File 1.

Of note were two mce-associated genes thought to be involved in macrophage invasion (Gagneux 2018) that showed MODERATE or MODIFIER impact annotations: *mce1C* (missense variant) and *mce1B* (downstream gene variant). A MODERATE impact variant (missense variant) was also flagged in *zmp1*, which may be involved in host-pathogen interaction (Correa et al. 2014).

Additionally, there were numerous variant positions that had MODERATE or MODIFIER impact annotations in genes involved in cell wall or membrane functions, as determined through Mycobrowser (Kapopoulou et al. 2011): *mmpL4* (missense variant), *mmpS4* (downstream gene variant), Rv0102 (upstream gene variant), Rv0912 (downstream gene variant), *ctpG* (missense variant), Rv1999c (missense variant), *cysA1* (missense variant), *cysW* (missense variant), *lppR* (downstream gene variant), *merT* (missense variant), Rv3104c (downstream gene variant), Rv3273 (missense variant), Rv3635 (downstream gene variant), and Rv3821 (downstream gene variant).

**Birth-death model parameterization**

For both datasets, birth death multi-rho tree priors were utilized. The L4 birth death skyline analyses were parameterized as follows for both BDSKY+UCLD+origin and BDSKY+UCLD. The rho parameter, referencing the sampling proportion at each sampling time, was split into four dimensions. One dimension was provided for each ancient sample, with a Beta prior distribution with mean 0.01, and one dimension was provided for modern genome representation, with Beta prior distribution with mean 0.1. The rho sampling times for each ancient genome were given as their respective tip dates, and the rho sampling time for modern genomes was set to 0. The reproductive number parameter (R), was given 5 dimensions, which provided estimates of population dynamics within L4 over time. The becomeUninfectiousRate was given one dimension for the ancient data with a Beta prior distribution with mean 0.001, and one dimension for the modern data with a Beta prior distribution of 0.1. For the L4 BDSKY+UCLD+origin analysis we imposed an upper limit on the origin of 10,000 years, with a starting value of 4,000 years (the initial value of the origin must be greater than the initial tree height estimate), and a uniform distribution.

MTBC birth-death analysis (BDSKY+UCLD) was parameterized as follows. The rho parameter was split into seven dimensions with one for each ancient genome, with a Beta prior distribution centered around 0.01 for dimensions one to six, and 0.1 for the seventh dimension, translating to 1% and 10% sampling probabilities for ancient and modern genomes, respectively. The sampling times for rho were set as the tip date for each ancient genome, and 0 for modern genomes. R was set to five dimensions. The becomeUninfectiousRate was given one dimension for the ancient data with a Beta prior distribution with mean 0.001, and one dimension for the modern data with a Beta prior distribution of 0.1.

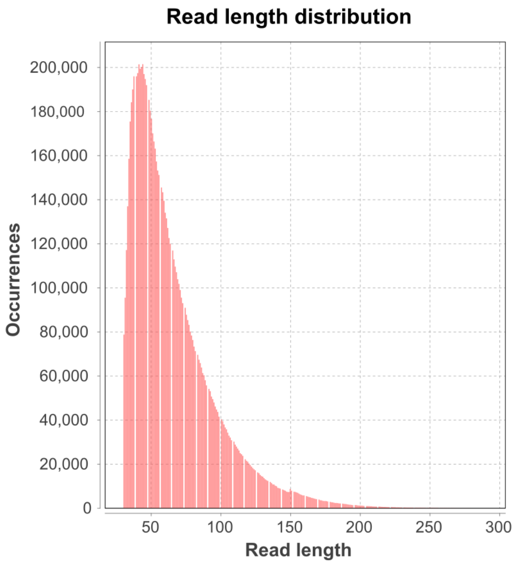


**Figure S1. Fragment length distribution plot for 150 cycle, paired-end sequencing data from UDG-treated, TB captured LUND1 library.** Fragment lengths as provided by DamageProfiler in the EAGER pipeline for 9,482,901 reads mapped to the TB ancestor genome with BWA as implemented in EAGER (-l 32, -n 0.1, -q 37).

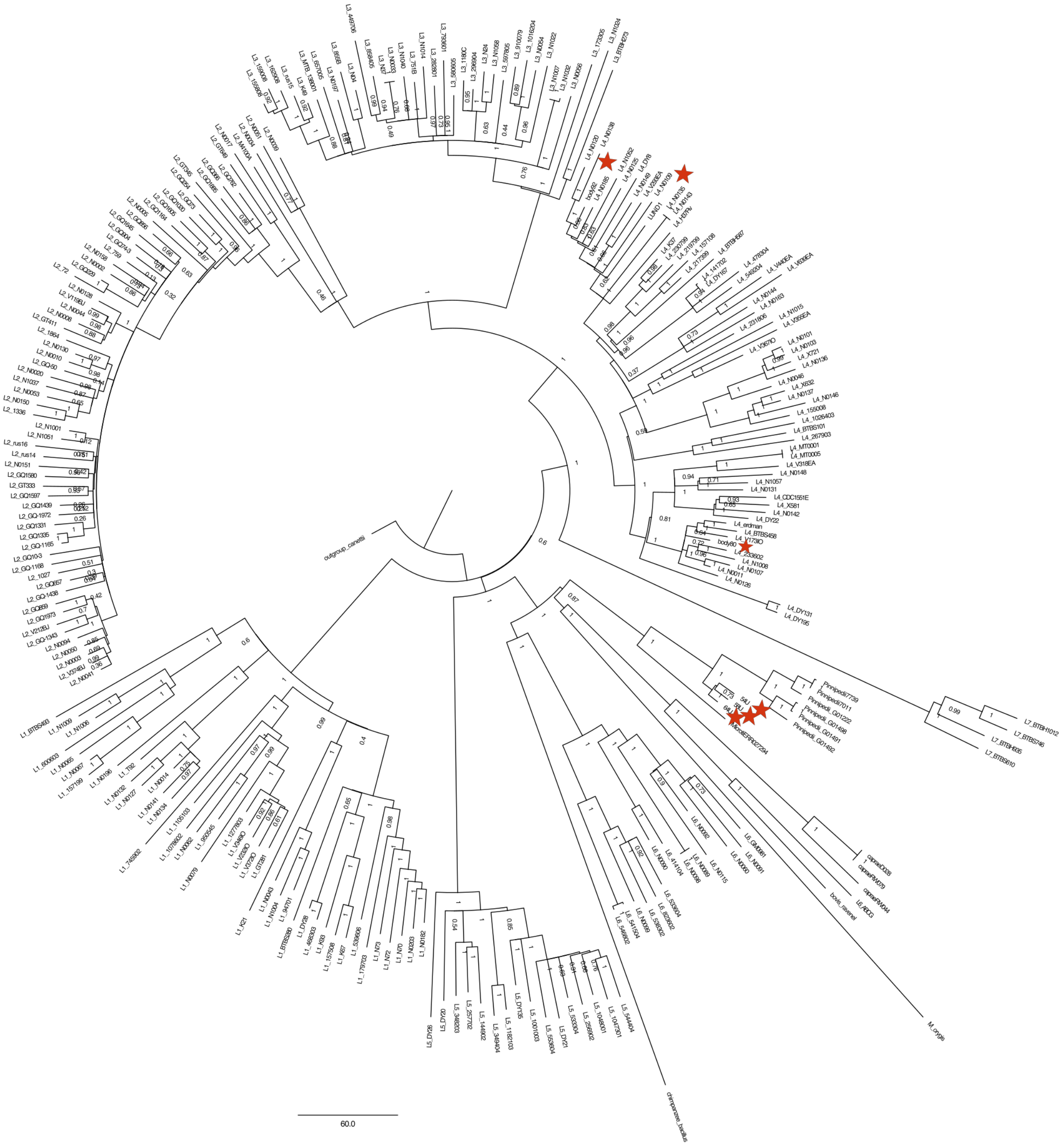

**Figure S2. MTBC dataset neighbor joining tree.** The neighbor joining tree was configured using MEGA-Proto and generated with MEGA-CC, with 500 bootstrap replicates. The red stars indicate the ancient genomes.

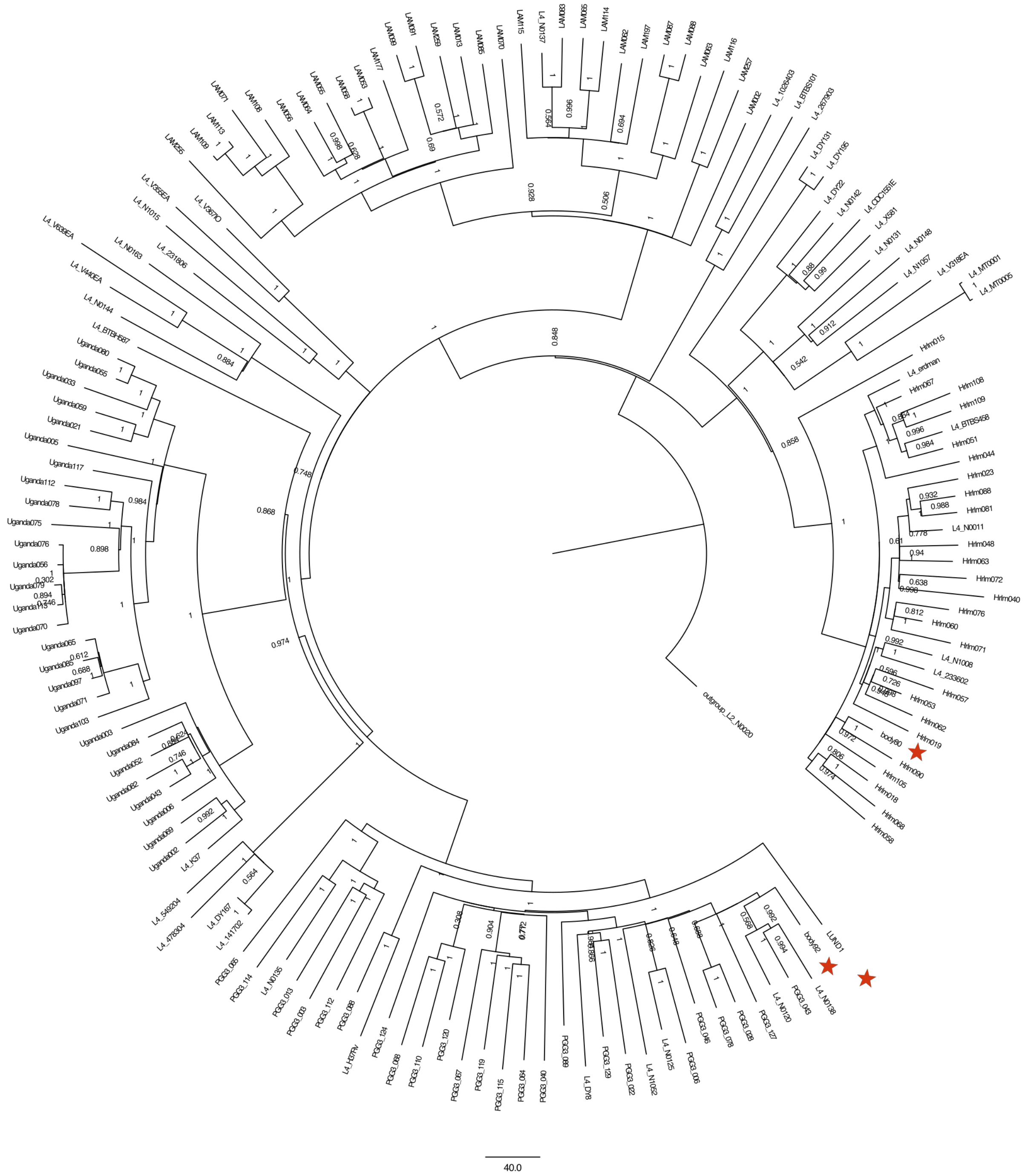

**Figure S3. L4 dataset neighbor joining tree.** The neighbor joining tree was configured using MEGA-Proto and generated with MEGA-CC, with 500 bootstrap replicates. The red stars indicate the ancient genomes.

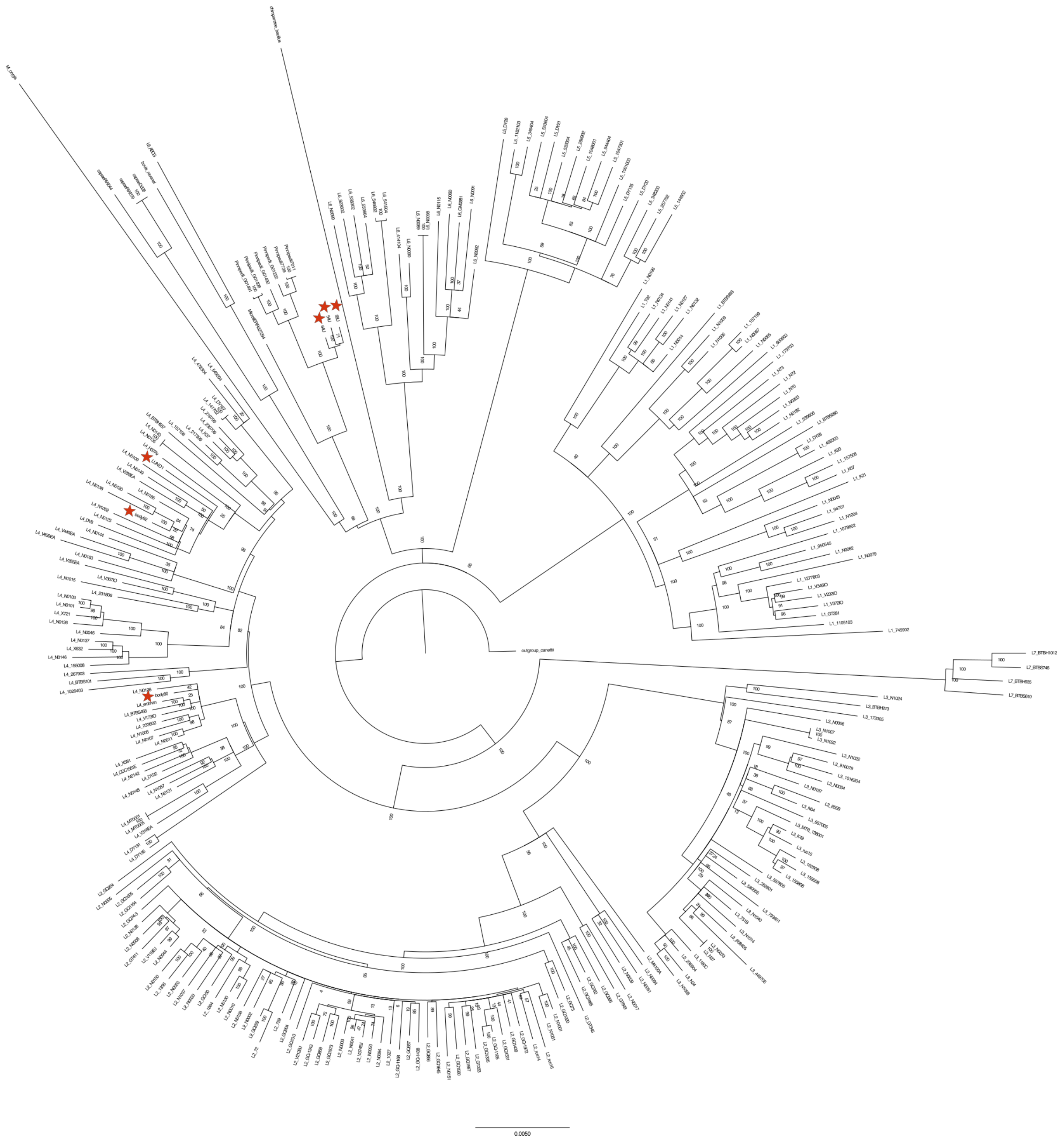

**Figure S4. MTBC dataset maximum likelihood tree.** The maximum likelihood tree was configured and generated with RAxML, with 500 bootstrap replicates. The red stars indicate the ancient genomes.

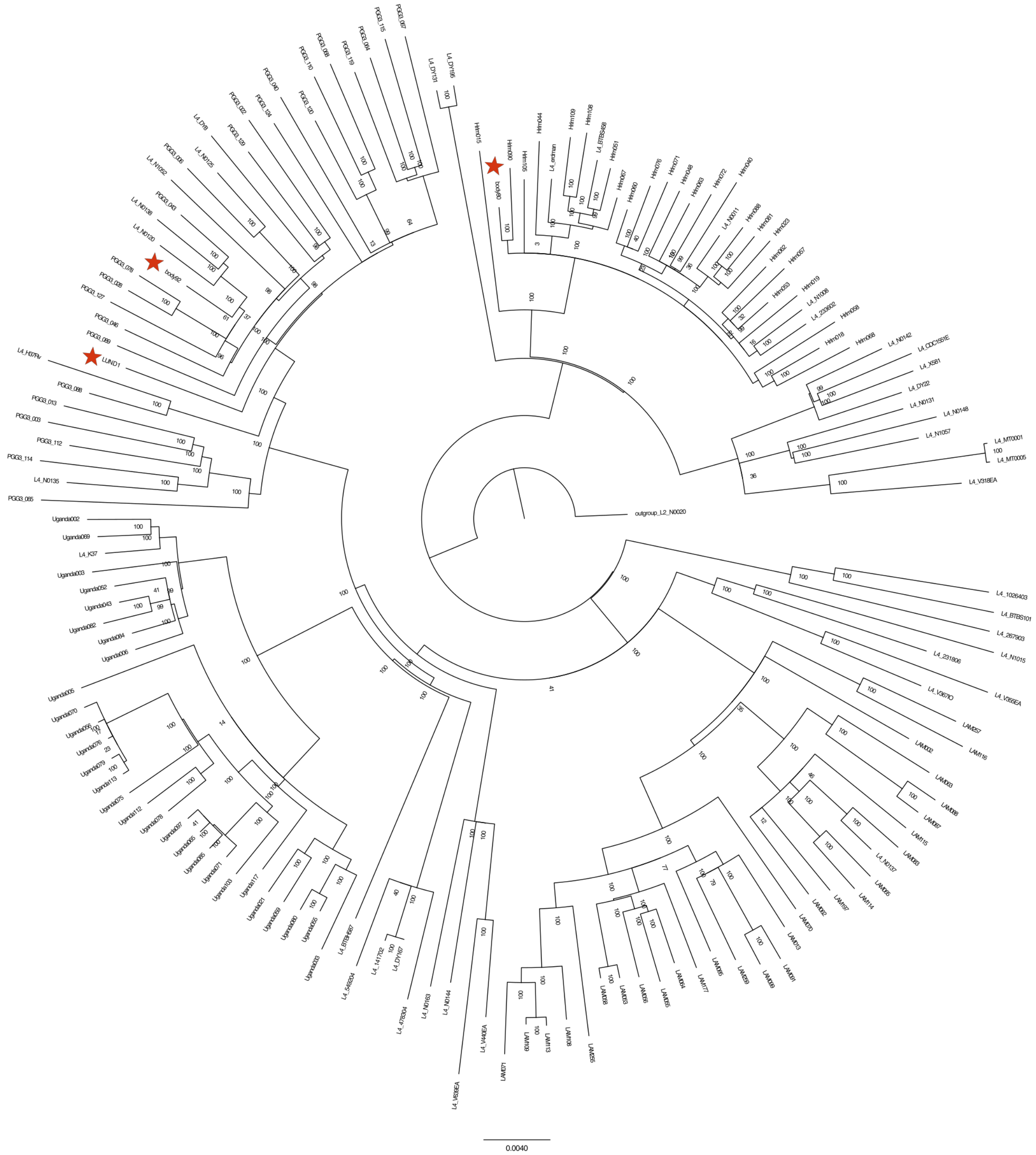


**Figure S5. L4 dataset maximum likelihood tree.** The maximum likelihood tree was configured and generated with RAxML, with 500 bootstrap replicates. The red stars indicate the ancient genomes.

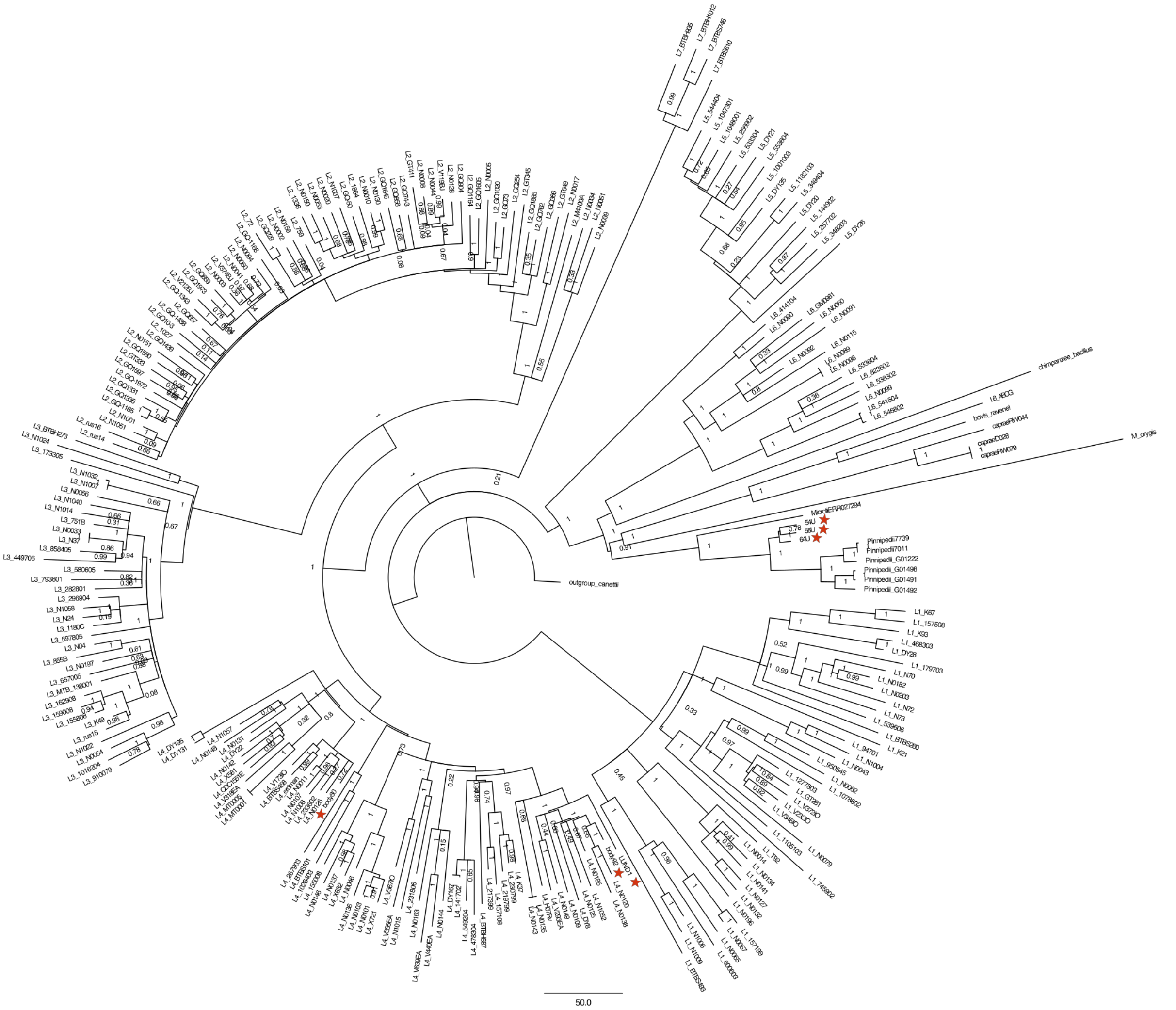

**Figure S6. MTBC dataset maximum parsimony tree.** The maximum parsimony tree was configured using MEGA-Proto and generated with MEGA-CC, with 500 bootstrap replicates. The red stars indicate the ancient genomes.

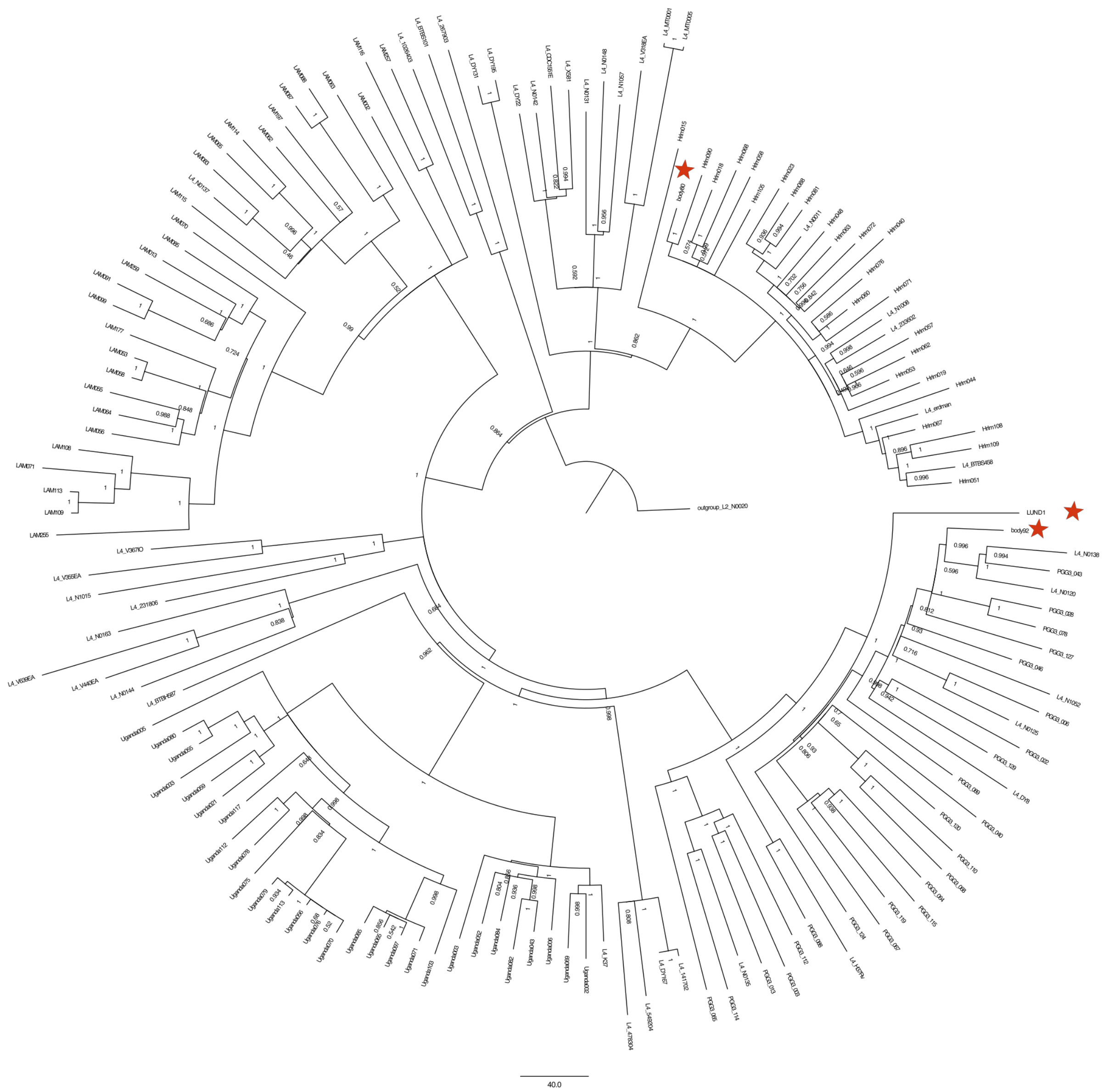

**Figure S7. L4 dataset maximum parsimony tree.** The maximum parsimony tree was configured using MEGA-Proto and generated with MEGA-CC, with 500 bootstrap replicates. The red stars indicate the ancient genomes.

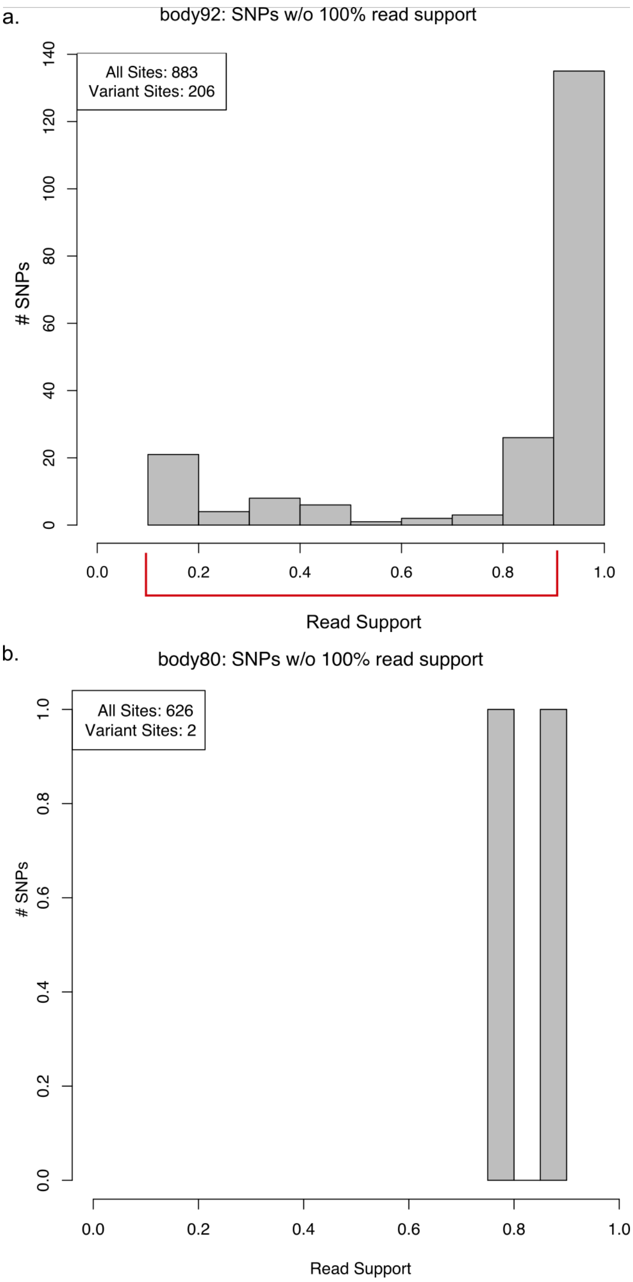


**Figure S8. Heterozygosity plots for body92 and body80.** A) The grey bars represent the quantity of sites at which between 10-99% of the reads represent an alternative derived allele (the allele must have coverage of 5-fold or greater to be counted). Body92 has 206 of these sites in total, though most of them have >90% representation, making them dominant alleles. The minority alleles, falling between 10-90% of the reads representing a given site (within the red bracket), are 70 in total. B) The grey bars represent the quantity of sites at which between 10-99% of the reads represent an alternative derived allele. Body80 has only two variant positions.

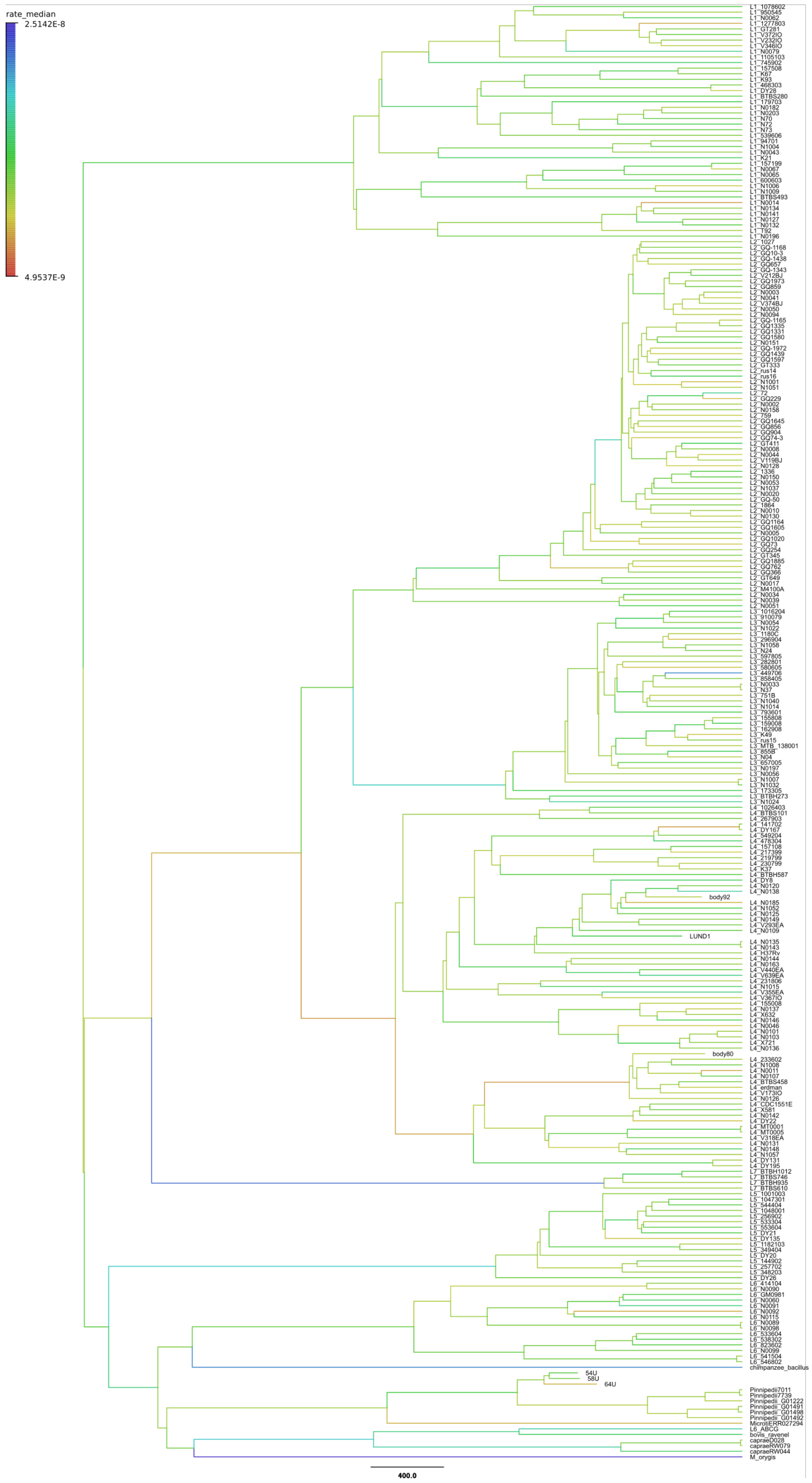


**Figure S9. MTBC BDSKY+UCLD maximum clade credibility tree.** Branches of the tree were colored according to median substitution rate.

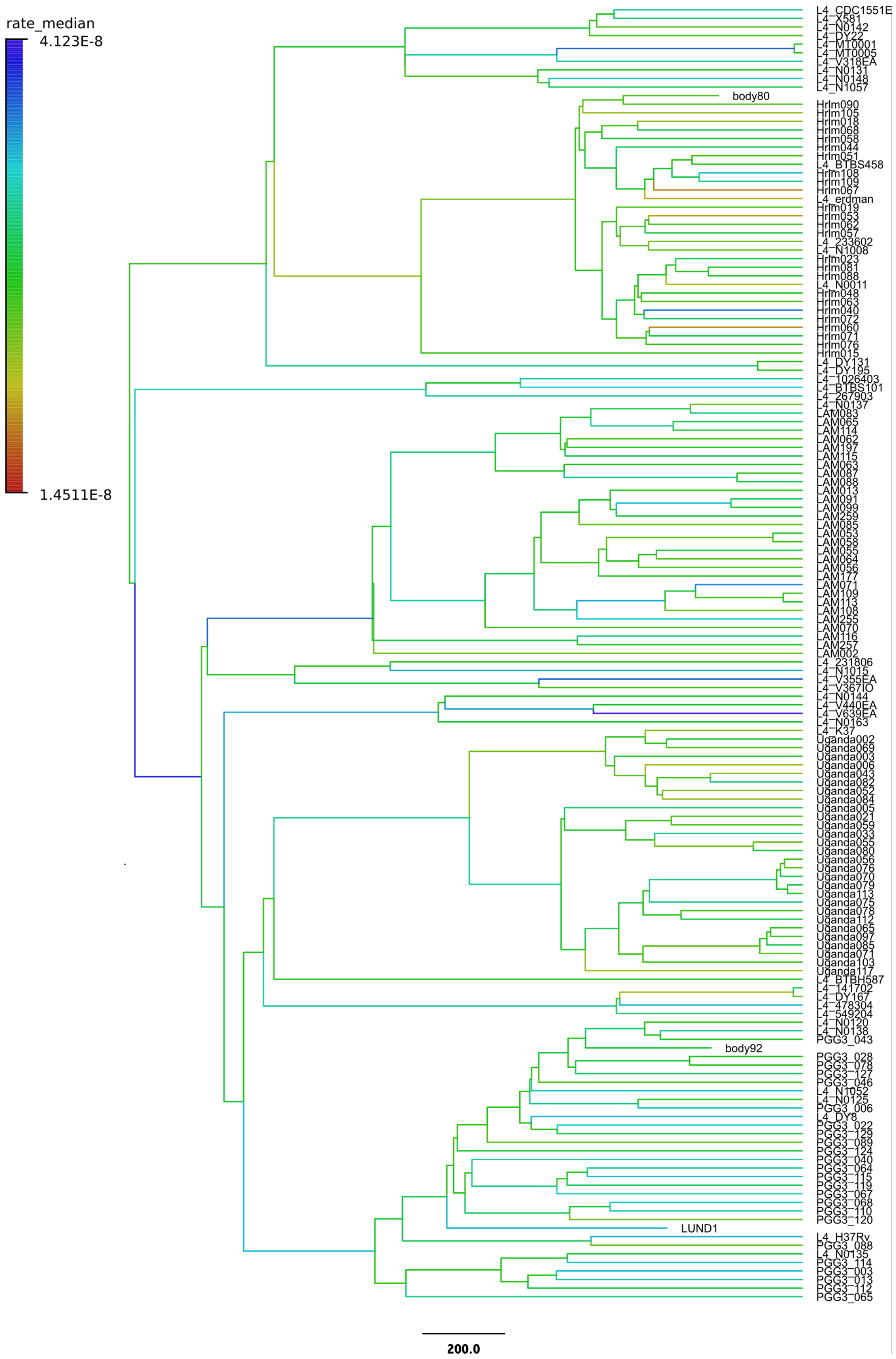


**Figure S10. L4 BDSKY+UCLD maximum clade credibility tree.** Branches of the tree were colored according to median substitution rate.
